# Epilepsy-linked kinase CDKL5 phosphorylates voltage-gated calcium channel Cav2.3, altering inactivation kinetics and neuronal excitability

**DOI:** 10.1101/2022.11.24.517538

**Authors:** Marisol Sampedro-Castañeda, Lucas L. Baltussen, Andre T. Lopes, Yichen Qiu, Liina Sirvio, Simeon R. Mihaylov, Suzanne Claxton, Jill C. Richardson, Gabriele Lignani, Sila Ultanir

## Abstract

Developmental and epileptic encephalopathies (DEEs) are a group of rare childhood disorders characterized by severe epilepsy and cognitive deficits. Numerous DEE genes have been discovered thanks to advances in genomic diagnosis, yet putative molecular links between these disorders are unknown. CDKL5 deficiency disorder (CDD, DEE2), one of the most common genetic epilepsies, is caused by loss-of-function mutations in the brain-enriched kinase CDKL5. To elucidate CDKL5 function, we looked for CDKL5 substrates using a SILAC-based phosphoproteomic screen. We identified the voltage-gated Ca^2+^ channel Cav2.3 (encoded by *CACNA1E*) as a novel physiological target of CDKL5 in mice and humans. Recombinant channel electrophysiology and interdisciplinary characterization of Cav2.3 phosphomutant mice revealed that loss of Cav2.3 phosphorylation leads to channel gain-of-function via slower inactivation and enhanced cholinergic stimulation, resulting in increased neuronal excitability. Our results thus show that CDD is partly a channelopathy. The properties of unphosphorylated Cav2.3 closely resemble those described for *CACNA1E* gain-of-function mutations causing DEE69, a disorder sharing clinical features with CDD. We show that these two single-gene diseases are mechanistically related and could be ameliorated with Cav2.3 inhibitors.

## Introduction

Developmental and epileptic encephalopathies (DEE) are characterized by severe early-onset epileptic activity accompanied by global developmental and cognitive impairments. The genetic aetiology approximately 90 DEEs has been identified ^1^, but targeted therapies remain scarce. Elucidating shared pathways in infantile-onset epilepsies will reveal molecular links between disease genes and greatly advance common targeted treatments.

<u>C</u>yclin <u>d</u>ependent <u>k</u>inase like-5 (CDKL5) is a brain-enriched serine-threonine kinase. *De novo* loss-of-function mutations in the X-linked *CDKL5* gene, including missense, nonsense and insertions/deletions, cause CDKL5 deficiency disorder (CDD) (OMIM 300672, 300203) ^2–5^. Pathogenic missense mutations, almost exclusively located in the kinase domain, indicate that kinase activity is critical for CDD pathology ^6–9^. CDD is characterized by infantile-onset, intractable seizures, profound neurodevelopmental impairment in motor and sensory function, impaired language acquisition and autonomic disturbances ^3, 5^. The estimated incidence for CDD is 1/42,000 live births each year ^10–12^ with approximately 80% being female patients, making it one of the most common types of genetic childhood epilepsy ^13, 14^.

CDKL5 is highly enriched throughout the forebrain with expression starting at late embryonic stages in rodents and humans ^15–19^. Low levels of expression were reported in other tissues ^16^. Phosphorylation targets include microtubule binding proteins EB2 and MAP1S ^6, 20^ and transcriptional regulators ELOA ^21^ and Sox9 ^22^, showing its diverse cellular roles. CDKL5 knockout (KO) neurons have deficits in synaptic transmission, dendritic spine development, neurite outgrowth, cilia elongation and microtubule dynamics ^23–29^. However, molecular mechanisms linking CDKL5 loss to neuronal hyperexcitability remain unknown. The identification of key physiological targets of CDKL5 directly involved in the regulation of cellular excitability may help elucidate epileptogenic mechanisms and lead to effective therapies.

Voltage-gated calcium channels (VGCC) have a key role in physiology, driving neuronal excitability and allowing influx of second messenger Ca^2+^ ions in response to membrane depolarization ^30–32^. VGCC subtype Cav2.3 is formed by the ion-conducting α1E subunit (encoded by *CACNA1E*) in complex with accessory subunits from the β and α2δ families. Cav2.3 is highly expressed in the CNS, where it is localized at both pre-and post-synaptic sites in neurons, as shown functionally and by immuno-EM ^33–35^. *CACNA1E* knockout mice demonstrated that Cav2.3 mediates neuronal R-type Ca^+2^ currents ^35–37^, so-called due to their “resistance” to organic VGCC channel inhibitors ^38, 39^. Cav2.3 is dynamically regulated by phosphorylation at multiple sites ^40, 41^. In general, VGCC phosphorylation is an important mechanism by which neuromodulators and their G-protein coupled receptors (GPCRs) exert their effect on cellular excitability ^42^.

*De novo* heterozygous missense mutations in *CACNA1E* cause an infantile epileptic encephalopathy (OMIM 618285) ^43, 44^. Cav2.3 variants studied *in vitro* have altered gating and/or inactivation kinetics, leading to gain-of-function (GoF) of Cav2.3 ^43, 44^. Patients with *CACNA1E* mutations have overlapping clinical phenotypes with CDD, such as intractable seizures, profound intellectual disability and hypotonia. Single-gene DEEs such as this one are estimated to impact approximately 1/2100 live births ^10^, but their real prevalence is unknown as more genes are being identified rapidly due to recent advances in genetic diagnosis ^45, 46^. Despite, overlapping clinical features, potential molecular links between these rare disorders remain to be elucidated.

In this work, we apply for the first time a global phosphoproteomics approach in CDKL5 knockout mice and identify and validate Cav2.3 as a novel substrate of CDKL5 in human and mouse neurons. Through our analysis of Cav2.3 function and characterization of a novel Cav2.3 phosphomutant mouse line, in which CDKL5 phosphorylation site is mutated, we reveal that lack of phosphorylation leads to Cav2.3 gain-of-function and hyperexcitability. Our results indicate that developmental and epileptic encephalopathies caused by *CDKL5* and *CACNA1E* are related at the molecular level. We propose Cav2.3 as a converging therapeutic target for these disorders.

## Results

### Cav2.3 is a CDKL5 phosphorylation substrate in mice and humans

We used stable isotope labelling of amino acids in cell culture (SILAC) ^47^ to differentially tag newly synthesized proteins in primary cortical cultures from wild type (WT) and CDKL5 knockout (KO) mice ^48^ and compare changes in protein phosphorylation levels (Fig. 1a). As expected in CDKL5 KO neurons, we observe a significant reduction in CDKL5 pS407 and in EB2 pS222, a known CDKL5 phosphorylation target ^20^. Phosphorylation of EB2 S221 was also decreased, indicating co-regulation of these neighbouring sites. More importantly, a potential new target protein, the ion channel Cav2.3, showed strongly reduced phosphorylation at S15 (pS15), in CDKL5 KO neurons. Strikingly, the Cav2.3 S15 site matches the RPXS/T* consensus motif ^6, 20^, suggesting direct CDKL5 phosphorylation (Fig. 1a, b). S15, located in the N-terminus of the α1E channel pore subunit (Fig. 1b), is conserved in humans (S14) but absent in the related Cav2.1 (*CACNA1A*) and Cav2.2 (*CACNA1B*) channels (Fig. 1b).

**Fig. 1:**
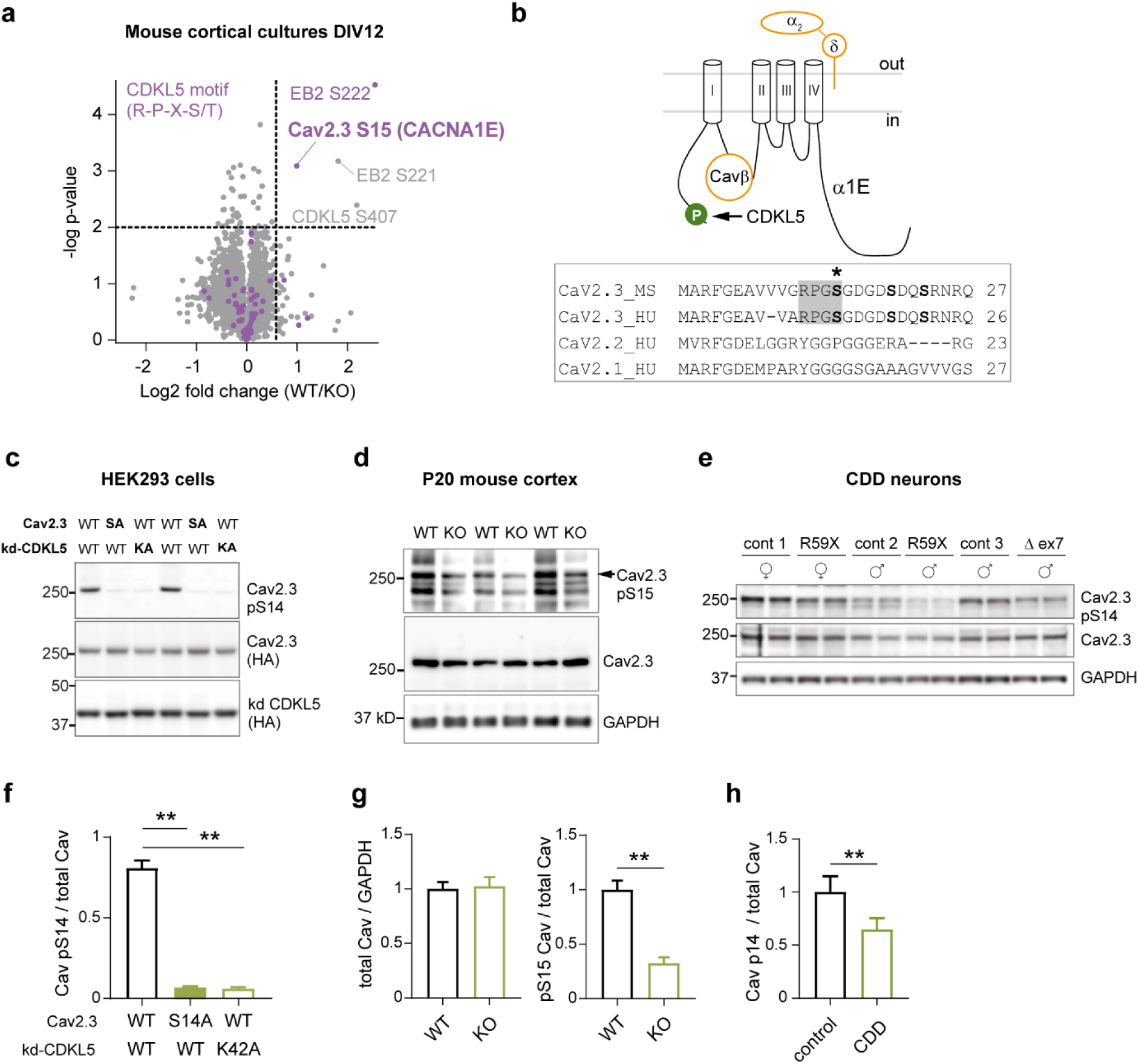
Identification and validation of Cav2.3 as substrate of CDKL5 kinase. **a** Volcano plot of differential phosphoprotein levels between WT and CDKL5 KO primary neurons, obtained using SILAC based quantitative proteomics (5 embryos/genotype). Each point represents one peptide; those with CDKL5 consensus motif RPXS/T are highlighted in magenta. The x axis shows log2 transformed fold change in phosphopeptide levels between genotypes and the y axis shows significance as -log10 transformed p value. Significance was established as indicated by the dotted lines (p<0.01, fold change 1.5, one sample t test). **b** Top: Schematic depiction of Cav2.3 formed of α1E channel pore subunit and associated proteins Cavβ and α2δ. CDKL5 phosphorylation site at α1E S15/14 is highlighted in green. Bottom: Alignment of proximal N-terminus of mouse and human Cav2.3 and related human Cav2.1 and 2.2 channel proteins. *S is the site of CDKL5 phosphorylation. **c** Western blot of HEK293 cells stably expressing human β3/α2δ1 subunits and transiently co-transfected with human HA-α1E (Cav2.3: WT or S14A mutant) and HA-CDKL5 kinase domain (kd CDKL5: WT or K42A mutant). S14 phosphorylation (pS14 Cav2.3) was detected with a custom made phosphoantibody. **d** Western blot of P20 cortices from WT and CDKL5 KO mice. **e** Western blot of iPSC-derived forebrain neurons from three CDD patients and related controls (parents) at 6 weeks of differentiation. Gender and *CDKL5* mutation are specified ^83^. **f** Quantification for immunoblot in **c** (**p<0.0001 Kruskal-Wallis ANOVA & Dunn’s test, n=11, 4 blots, 2 transfections). **g** Quantification for immunoblot in **d** (**p<0.0001 two tailed unpaired t test, n=9, 3 blots, 6 mice/genotype). **h** Quantification for WB in **e** (**p<0.005 two tailed paired t test, n=6, 2 blots, 2 technical replicates). For antibodies used see Methods.

To test if CDKL5 can phosphorylate Cav2.3 S15, we generated a phosphospecific antibody targeting this site. Overexpression of Cav2.3 (& ancillary subunits) together with the CDKL5 kinase domain in HEK293 cells shows strong Cav2.3 phosphorylation *in vitro* (Cav2.3 pS14, Fig. 1c, f). This phosphorylation is absent when overexpressing phosphomutant Cav2.3 or kinase-dead CDKL5, demonstrating the specificity of the antibody and the dependence of phosphorylation at this site on CDKL5 kinase activity. Importantly, full-length CDKL5 also directly phosphorylated Cav2.3 *in vitro* (Supplementary Fig. 1a, b). Cav2.3 phosphorylation is consistently reduced in the cortex of CDKL5 KO mice at postnatal day P20 (Cav2.3 pS15), validating the substrate *in vivo* (Fig. 1d, g). Finally, Cav2.3 pS14 was decreased in human iPSC-derived neurons from CDD patients (Fig. 1e, h), demonstrating altered phospho-regulation of this target in CDD. Together, these results show that Cav2.3 is a *bona fide* CDKL5 phosphorylation target in mice and humans.

### Loss of Cav2.3 Ser14 phosphorylation slows baseline channel kinetics and amplifies muscarinic regulation

Cav2.3 is causally linked to epilepsy by two main lines of evidence. First, channel deletion renders mice resistant to tonic-clonic seizures ^49–51^. On the other hand, Cav2.3 GoF leads to severe epileptic encephalopathy in humans ^43, 44^. The remarkable overlap in clinical manifestations between CACNA1E and CDD patients, including seizures and neurodevelopmental delays, raises the possibility that Cav2.3 regulation by CDKL5 might affect channel function and in turn neuronal excitability.

To test the effect of pS14 on Cav2.3 channel properties we used a HEK293 cell line stably expressing the human auxiliary subunits β3 and α2δ1 and transiently transfected human α1E and full-length CDKL5 plasmids (Supplementary Fig. 1a, b). Whole-cell patch-clamp recordings of Cav2.3 Ba^2+^ currents revealed slower decay kinetics (inactivation τ) in S14A mutant channels (Fig. 2a-c). Notably, in absence of CDKL5, WT Cav2.3 decay kinetics were indistinguishable from S14A Cav2.3 mutant (Fig. 2d). We also found a statistically significant increase in whole-cell current density at maximum activation voltages in S14A when compared to WT Cav2.3 in presence of CDKL5 (Fig. 2e). To account for differences in biophysical properties that may be introduced by α1E association with a different β subunit ^52, 53^, we conducted parallel experiments in a HEK293 cell line stably expressing human α2δ1 and β1b, another Cavβ subunit associated with α1E in the brain ^54^. We used Ca^2+^ as charge carrier to examine channel properties in a more physiological context. Decay kinetics of WT Cav2.3 currents with β1b were substantially slower than currents with β3 (Supplementary Fig. 2a, b *vs.* Fig 1a, b), as previously reported ^52^. In absence of CDKL5 phosphorylation (S14A α1E mutant or kinase-dead CDKL5 conditions), Cav2.3 current inactivation was further slowed (Supplementary Fig. 2a-c), recapitulating our results with β3 (Fig. 2a-c). Current density at maximal activation voltages did not reach significance in the β1-expressing cells, where currents were generally larger (Supplementary Fig. 2d). Finally, we observed no changes in half maximal activation and inactivation potential or time course of recovery from fast inactivation in β3-or β1-expressing cells (Fig. 2f and Supplementary 2e,f).

**Fig. 2:**
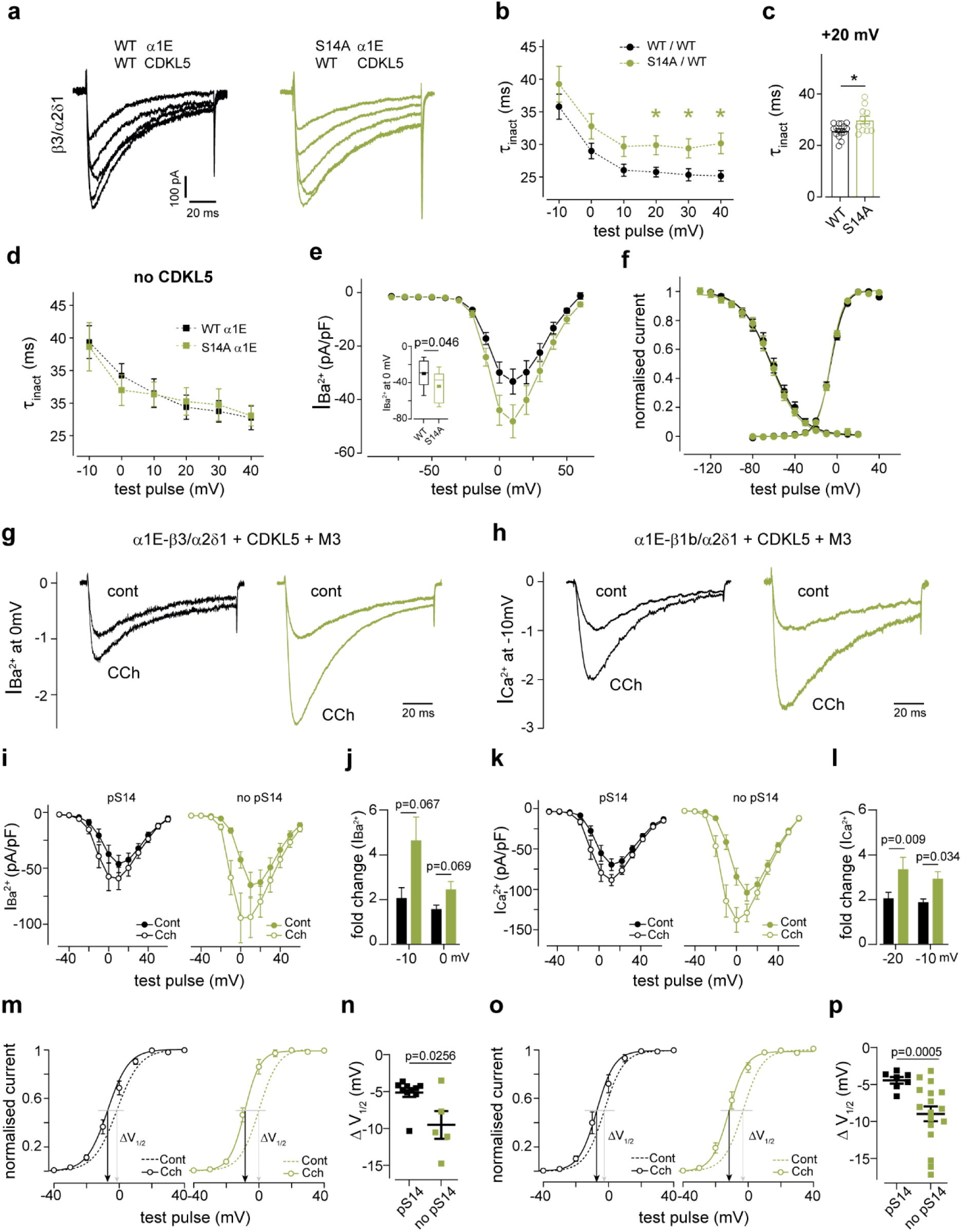
Functional characterization of phospho-Ser14 Cav2.3 in HEK 293 cells. **a** Depolarization-evoked current responses in HEK293 cells stably expressing human β3/α2δ1 subunits and co-transfected with human Cav2.3 (WT α1E or S14A α1E) and FLAG-CDKL5 full length (WT CDKL5). Colours denote different construct combinations. Traces show steps from -10 to +30mV from -80mV; Ba^2+^ was charge carrier b Open channel inactivation tau (τ_inact_) for Cav2.3 with (WT/WT, n=13-15) and without (S14A/WT, n=11-13) CDKL5 phosphorylation (*p=0.02, 0.03 & 0.01 at +10, +20 & +40 mV Two-Way ANOVA, Fisher’s LSD). Data were acquired using 100ms voltage steps from -80mV in +10mV increments every 10s. c Individual data points for τ_inact_ at +20mV (*p=0.01, two-tailed unpaired t test). d WT or phophomutant S14A Cav2.3 inactivation in the same cell line in absence of CDKL5 (n=7-9; p>0.05 Two-Way ANOVA); Ca^2+^ was charge carrier. e Current voltage relationship for cells in **b**. Inset: comparison of current density at 0mV (inset, two-tailed unpaired t test). **f** Normalised Cav2.3 current conductance and voltage dependence of inactivation for the same transfection conditions. Activation V_1/2,_ n: WT/WT -6±1 mV, 15; S14A/WT -6±1 mV,12; Inactivation V_1/2_, n: WT/WT -61±3 mV,10; S14A/WT -62±2 mV,11 (p>0.05, two tailed unpaired t tests). For inactivation protocol see Methods. Solid lines are Boltzman fits to the average data points **g** Normalised Ba^2+^ currents in the b3/a2d1-cell line or **h** Ca^2+^ currents in the b1/a2d1-cell line, co-transfected with α1E (WT or S14A), WT CDKL5 and muscarinic type 3 receptor (M3), in presence and absence of carbachol (CCh) 30 and 10 μM, respectively. HP= -100mV. **i** IV curves and **j** CCh-induced change in current amplitude for experiment in **g** (pS14 n=8, no pS14 n=7, Two-Way ANOVA, Fisher’s LSD). **k** IV curves and **l** CCh-induced change in current amplitude for experiment in **h** (pS14 n=7, no pS14 n=16, TW ANOVA, Fisher’s LSD). **m** Conductance plots for experiment in **g** and **n** Corresponding change in activation (V_1/2_: -5±0.6 *vs*. -10±1.9 mV, Kolmogorov-Smirnov). **o** Conductance plots for experiment in **h** and **p** corresponding change in activation (V_1/2_: -4±0.5 *vs* -9±1 mV, Welch’s t test). From **l-p**, ‘pS14’ refers to WT a1E/WT CDKL5; ‘no pS14’ refers to S14A α1E /WT CDKL5 and α1E WT /K42R* CDKL5 conditions pulled together.

Altogether, our results indicate that CDKL5-mediated Ser14 phosphorylation of α1E speeds up the inactivation of human Cav2.3 and suppresses current amplitude, independent of Cavβ subtype or charge carrier. Consequently, absence of CDKL5 leads to Cav2.3 GoF, with larger and more prolonged currents. A subset of *CACNA1E* GoF mutations described in human DEE patients also cause slower inactivation kinetics and/or increased current density ^43^, mirroring the phenotypes observed in the unphosphorylated channel. Thus, the functional deficits in Cav2.3 observed in absence of CDKL5 phosphorylation match those caused by pathological point mutations in Cav2.3 and could partly explain the pathophysiology of CDD.

Cav2.3 channels are downstream targets of muscarinic acetylcholine receptors, which modify channel function predominantly via PKC-mediated phosphorylation, leading to current enhancement in HEK293 cells and neurons ^55–57^. Given the observed GoF in S14A Cav2.3, we hypothesized that muscarinic receptor enhancement may also be affected by CDKL5 phosphorylation. To test this, we co-expressed Cav2.3, CDKL5 and muscarinic acetylcholine receptor type 3 or 1 (M3 or M1), which are prominent brain subtypes known to regulate neuronal Cav2.3 post-synaptically. As expected, application of the non-hydrolysable agonist carbachol substantially increased current amplitude regardless of Ser14 phosphorylation in both β3 and β1 stable cell lines (Fig. 2g-l). The current increase with carbachol was 1.5 to 2 fold greater in cells without phospho-Ser14 at -10 mV (Fig. 2g, h, j, l), reflecting Cav2.3 gain-of-function in absence of phosphorylation. The half-maximal activation potential (V_1/2_) was left-shifted by 5 mV for WT channels in both cell lines, a described effect of muscarinic receptor activation ^56, 58^. Interestingly, without Ser14 phosphorylation this shift was significantly greater, implying an effect of phospho-Ser14 on the modulation of Cav2.3 gating (Fig. 2m-p). This represents an additional gain-of-function due to loss of CDKL5, a phenotype comparable to pathogenic *CACNA1E* mutations that alter V_1/2_ ^43^.

Upon GPCR activation, Cav2.3 channels can be suppressed by G_βγ_ subunits. This effect is has been observed in Gαq-coupled receptors including M1/M3 in HEK293 cells ^56^ and CA1 neurons ^55^. The Cav2.3 N-terminus and Cavβ are involved in G_βγ_ regulation ^59, 60^. For this reason, we investigated the possibility that the enhanced carbachol effects in absence of N-terminal pS14 could be due to reduced G_βγ_ suppression. To distinguish between PKC-mediated enhancement and G_βγ_ suppression, we boosted PKC activity prior to carbachol application using phorbol 12-myristate 13-acetate (PMA), reliably increasing Cav2.3 currents ^56, 61^. With further addition of carbachol we found no evidence of muscarinic inhibition of human WT Cav2.3 channels (Supplementary Fig. 2g, h). Moreover, co-expression of Cav2.3 and Gi-coupled dopamine D2 receptors to further probe G_βγ_ inhibition in our system, revealed significant reduction in peak currents upon D2 activation with quinpirole ^59^. However, there was no difference in inhibitory modulation with or without Ser14 phosphorylation (Supplementary Fig. 2i, j). The above manipulations suggest that G_βγ_ is not involved in pathological Cav2.3 GoF.

Our results describe two Cav2.3 gain-of-function (GoF) mechanisms in the absence of CDKL5 phosphorylation: slower decay kinetics and a left shift in voltage-dependence of activation (V_1/2_) upon muscarinic activation, both reflected in increased Ca^2+^ influx through Cav2.3 channels. These GoF features are reminiscent of those described for pathological Cav2.3 variants.

### Altered R-type current inactivation and increased excitability in neurons from Cav2.3 S15A phosphomutant mice

To investigate the role of Cav2.3 S15 phosphorylation in neurons, we generated Cav2.3 S15A phosphomutant mice using CRISPR-Cas9 genome editing. We found that the total S15 phosphorylation levels detected by our phosphoantibody were reduced to <20% of control levels in homozygous S15A mice (HOM S15A) while total Cav2.3 levels were not changed (Fig. 3a-c). Cav2.3 is highly expressed in the somato-dendritic region of CA1 hippocampal neurons ^34^ where it underlies most of the R-type current and regulates intrinsic excitability ^35, 55^. We performed whole-cell somatic recordings in acute slices from young mice and recorded pharmacologically isolated Cav2.3 Ca^2+^ currents in these cells. When compared to wild type littermates (WT), HOM S15A mice showed significantly slower inactivation τ at maximal conductance voltages, recapitulating HEK293 cell results (Fig. 3d, e). Current amplitudes at the soma did not differ (Fig. 3f).

**Fig. 3:**
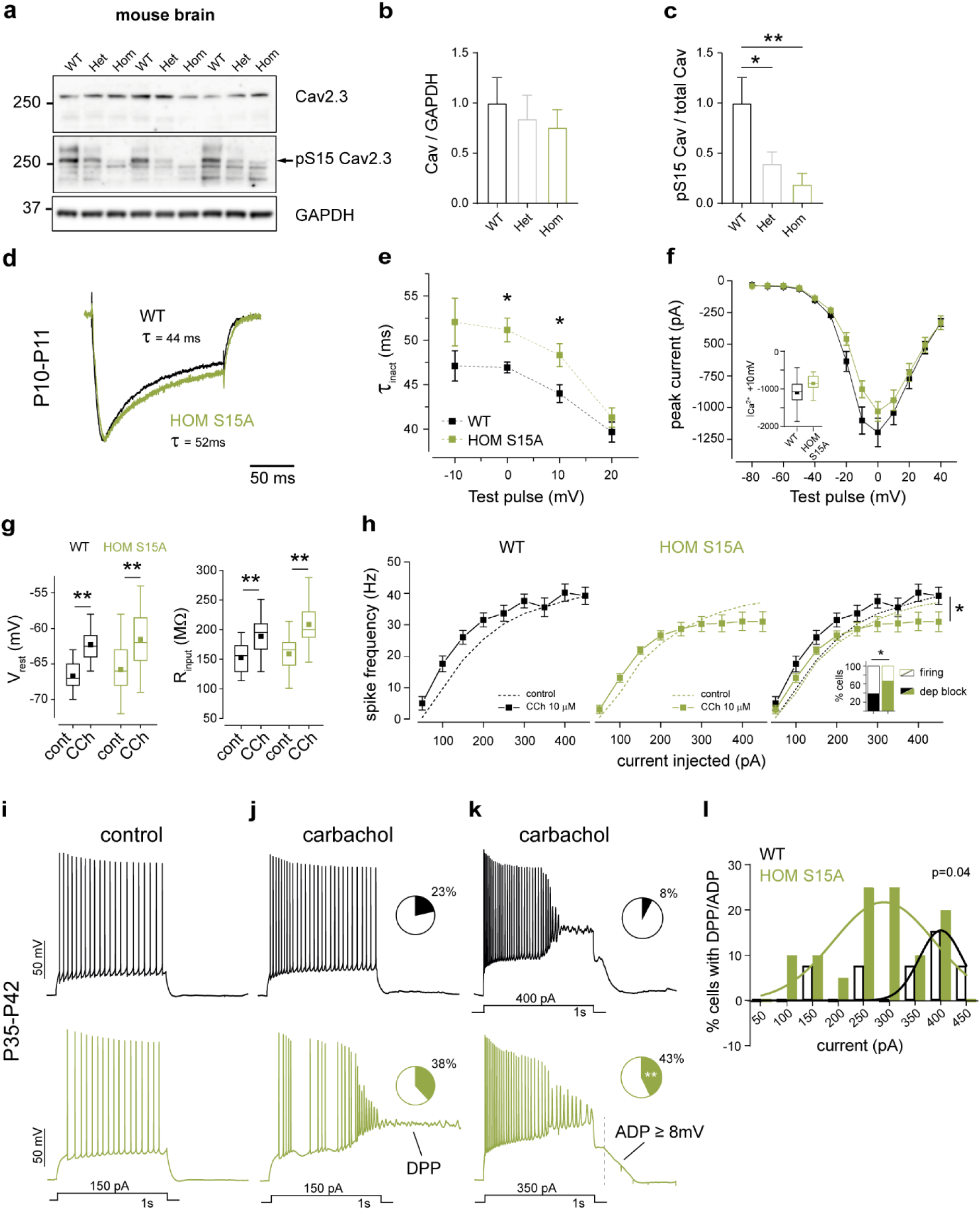
Physiological properties and cholinergic neuromodulation of hippocampal CA1 neurons from WT and phosphomutant mice. **a** WB validation of CRISPR-generated Cav2.3 Ser15Ala mutant mice using a pS15 phosphoantibody in whole-brain lysates from 12-week-old WT, Heterozygous S15A and Homozygous S15A mice. **b** Quantification of total Cav2.3 (p>0.05 One-Way ANOVA) and **c** relative phospho-S15 Cav2.3 (*p=0.029, **0.006 One-Way ANOVA, Fisher’s LSD) expression (n=5, 2 blots, 3 mice/genotype). **d** Normalised R-type Ca^2+^ current in neurons from young mice of each genotype during a step from -60 to 0 mV in presence of Na^+^, K^+^, Ca^2+^ and synaptic channel inhibitors. **e** R-type current inactivation tau (τ_inact_) at maximal activation voltages (*p=0.01 Two-Way Repeated Measures ANOVA, Fisher’s LSD) and **f** average IV curve for all P10-11cells recorded (p>0.05), (n=13 neurons, 4-5 mice/genotype). Data for -10mV step is shown in the inset (p=0.06, two-tailed unpaired t test). **g** Membrane resting potential (V_rest_, left) and input resistance (R_input_, right) of adult neurons (WT n=13, HOM S15A n=17; 7 mice/genotype) in control conditions and after 10μM carbachol (CCh) application (**p<0.002, two-tailed paired t test). **h** Input/output curves for the same neurons (WT n=9, HOM n=13, *p<0.028 Two-Way Repeated Measures ANOVA) and percentage of cells with CCh-induced depolarization block (inset, WT n=13, HOM n=21, *p<0.012, binomial test). **i** Representative control action potential trains evoked in WT (top) and HOM S15A (bottom) CA1 cells by 1s-long current injections. Membrane was held at - 65mV. **j** Same cells as **i** upon application of CCh and representative depolarising plateau potential (DPP), observed at some current injections in both genotypes; insets: fraction of cells with at least one DPP in each genotype group (WT n=13, HOM n=21, p>0.05 binomial test). **k** Same cells as **j** at higher stimulation illustrating sustained depolarization and AP block (top) or attenuation (bottom) towards the end of the stimulus and long-lasting large amplitude afterdepolarizations (ADP). Like DPPs, these ADPs were present at some current injections in a fraction of cells in both groups of mice (inset, WT n=13, HOM n=21, **p<0.004, binomial test). **l** Percentage of cells displaying sustained DPPs or ADPs upon stimulus termination at each current injection step (non-linear fit comparison).

We next tested the effect of Cav2.3 S15A on neuronal excitability and cholinergic neuromodulation in response to depolarising current injections in adult CA1 pyramidal cells. Resting membrane potential (V_rest_), input resistance (R_in_) and firing properties were identical between WT and S15A mice under baseline conditions (Fig. 3g-i). Following bath application of carbachol, we observed a 4 mV depolarisation in V_rest_ and a 25% increase in R_in_ (Fig 3g), changes that were comparable between WT and S15A neurons (Supplementary Fig. 3a). These alterations are likely due to conductances other than Cav2.3, in line with previous _reports 58, 62, 63._

In neurons, cholinergic activation can elicit sustained depolarizing plateau potentials (DPPs): all-or-none, large amplitude depolarizations that outlast stimulus-evoked firing ^58, 64, 65^. A muscarinic medium duration afterdepolarization (ADP) is also reported in CA1 pyramidal cells ^66^. Up-regulation of Cav2.3-mediated R-type currents is required for DPPs and ADPs in hippocampus ^67^ and cortex ^65^. We inspected these Cav2.3-dependent hyper-excitability measures in S15A mice. Carbachol application significantly increased action potential firing frequency in WT and S15A neurons (Fig. 3 h-k). At current injections greater than 250 pA, however, phosphomutant neurons responded with fewer spikes and depolarization block towards the end of the stimulus step, indicating greater underlying depolarization. The percentage of cells exhibiting at least one DPP or all-or-none afterdepolarization (ADP) >8 mV upon carbachol treatment was significantly higher in S15A mice, showing an enhanced depolarization response (Fig. 3j, k). Importantly, the occurrence of DPP/large ADPs was shifted towards lower current injections in S15A cells (Fig. 3l). Greater perisomatic depolarization in S15A neurons was also reflected in a larger ADP at 100ms post stimulus and more spike attenuation (Supplementary Fig. 3b, c). Together with the enhanced carbachol-mediated left-shift in V_1/2_ observed in phosphomutant channels in HEK293 cells (Fig. 2 j,l), these increased neuronal depolarizations in Cav2.3 phosphomutant mice suggest facilitation of gating of endogenous Cav2.3 without S15 phosphorylation upon cholinergic agonist application.

We tested if other conductances known to be regulated by carbachol are altered in Cav2.3 phosphomutants. The medium-and slow-afterhyperpolarizations (AHPs) are critical for spike frequency regulation and their underlying K^+^ and mixed-cationic currents are targets of the muscarinic system ^62, 68, 69^. Baseline AHPs elicited by a short spike burst were not different between WT and S15A mice and were equally suppressed by carbachol application (Supplementary Fig. 3d). We also analysed baseline small-conductance Ca^2+^-activated K^+^ (SK) currents and spontaneous excitatory synaptic currents (EPSCs), known to be functionally coupled to ^70^ or modulated by ^71^ Cav2.3 in CA1 neurons. We hypothesized that these may be affected given the prolonged Cav2.3 currents in S15A mice, but our measurements show no differences between genotypes (Supplementary Fig. 3e-h). Therefore, we suggest that the increased carbachol-induced excitability of CA1 neurons in Cav2.3 S15A knock-in mice is specifically due to Cav2.3 GoF.

### Cav2.3 S15A phosphomutant mice have behavioural and EEG deficits

Multiple CDKL5 KO mouse models have been generated by deleting exons 4, 2 or 6, all of which led to loss of CDKL5 protein ^15, 18, 48^. Numerous behavioural deficits were reported in CDKL5 KO mice ^15, 48, 72–74^, thus we investigated behavioural phenotypes in Cav2.3 S15A phosphomutant mice to see if any of the CDKL5 KO impairments are recapitulated. We observed no spontaneous behavioural seizures in Cav2.3 S15A, as reported for CDKL5 KO mice ^15, 18, 48^, but see ^75, 76^. When monitored in a home-cage environment for prolonged periods, S15A showed reduced locomotion and voluntary wheel use (Fig. 4a, b), also observed in CDKL5 KO mice ^48^. We observed minor changes in locomotion and zone preference in the open field test, but only towards the end of a 30 min trial (Supplementary Fig. 4c, 4a), potentially indicating fatigue rather than anxiety. Unlike CDKL5 KOs, S15A mice did not present hindlimb clasping (mean scores: WT 0.041, HOM S15A 0, p=0.334, unpaired t test) and performed equally in the rotarod test (Fig. 4d) when compared to WT littermates. Cognitive abilities in the Y-and Barnes maze were also unchanged in phosphomutants (Supplementary Fig. 4b-e). On the other hand, S15A mice had mild impairments in sociability (Fig. 4e). A consistently reported phenotype of CDKL5 KO mice is altered contextual fear conditioning responses ^15, 73^. Memory formation, retention and extinction in this task were also defective in Cav2.3 S15A (Fig. 4f), indicating a clear overlap with CDKL5 KOs.

**Fig. 4:**
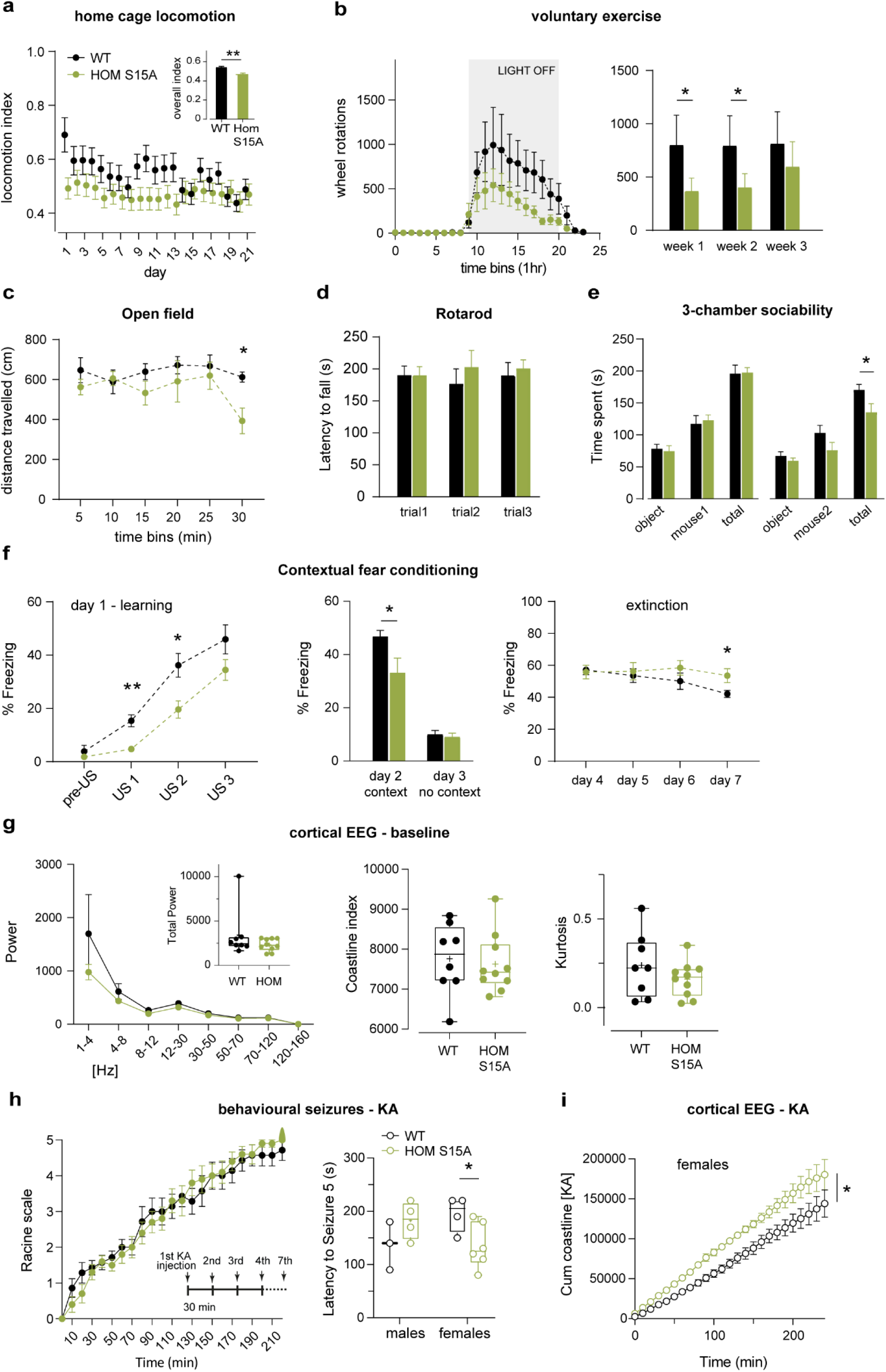
*In vivo* phenotypic characterization of Cav2.3 S15A mice. **a** Average home cage night-time locomotion of WT and HOM S15A mice (N=9) per day and overall locomotion index (inset) in a three-week period (**p<0.0001, Welch’s t test). **b** left: wheel rotations per hour during voluntary exercise activity for the same mouse cohort as **a**, over a two-week period; right: overall dark cycle average rotations in each group for three consecutive weeks (week 1 *p=0.017, week 2 *p=0.049 Two-Way ANOVA, Fisher’s LSD). **c** Total distance travelled by WT (N=8) and HOM S15A (n=7) mice in an open field arena (p<0.013 at 30 min Two-Way ANOVA, Fisher’s LSD). **d** Accelerating rotarod performance for the same cohort as **c** (p>0.05, TW ANOVA). Animal weights were equal (WT=25±4 g, HOM=25±5 g, p>0.05, two-tailed unpaired t test). **e** Social, non-social and total exploration times during a 10min three-chamber sociability test (*p=0.04, two-tailed unpaired t test). In habituation trials, chamber occupancy was equal between groups. **f** Freezing behaviour during learning day 1 (left, *p=0.01, **p=0.002), associative memory test day 2+3 (middle, *p=0.03) and memory extinction test days 4-7 (right, *p=0.043), in a fear-conditioning experiment with tactile and olfactory cues (Two-tailed unpaired t test or Two-Way Repeated Measures ANOVA, Fisher’s LSD). **g** left: two-week baseline ECoG spectral analysis and total power (inset) in WT (N=8) and HOMS15A (N=10) 9-12 week-old mice; corresponding coastline index (middle) and kurtosis (right) (p>0.05, Two-Way ANOVA and two-tailed unpaired t tests). No sex differences were observed. **h** left: behavioural scoring of kainic acid (KA)-induced seizures (WT N=7, HOMS15A N=10, p>0.05, Two-Way ANOVA) using repetitive 5mg/kg i.p. injections (inset); right: latency to stage 5 tonic clonic seizures per gender (males, p>0.05; females *p<0.0449, two-tailed unpaired t test). **i** Corresponding cumulative ECoG coastline during KA seizures in females (WT N=3, HOM S15A N=4, *p<0.018, Two-Way ANOVA).

Finally, given the high epilepsy incidence in human CDKL5 and CACNA1E DEE patients, seizure susceptibility was analysed in a separate cohort using wireless electrocorticogram (ECoG) transmitters ^77^ and repeated low-dose kainic-acid (KA) injections ^78^ (Fig. 4g-i). We found no changes in baseline ECoG (Fig 4g) or overall KA-induced seizure susceptibility in S15A mice (Fig. 4h), similar to KA-induced seizures in CDKL5 KO mice ^18, 48^. Interestingly, however, female S15A mice had a reduced threshold to stage 5 behavioural seizures with an increase in ECoG activity during KA (Fig. 4h,i).

In summary, our *in vivo* analyses of Cav2.3 S15A mice demonstrate similarities with CDD mouse models and patients, including social, motor and cognitive impairments, and to some extent increased seizure susceptibility. These observations suggest that loss of pS15 is a key contributor to the phenotypic features of CDD.

## Discussion

There are currently no disease-targeting therapies for CDD and the causes of aberrant excitability in absence of CDKL5 are unknown. Here we identify the alpha subunit of Cav2.3 channel complex as a CDKL5 phosphorylation substrate in mouse and human neurons. *CACNA1E* mutations cause severe early onset epilepsy in humans and there is remarkable overlap between CDD and patients with *CACNA1E* mutations. These include intractable seizures, global developmental delay, intellectual disability, hypotonia, hyperkinetic movements and sleep disturbances. A consistent feature of Cav2.3 variants associated with epilepsy is GoF of the channel, by left-shifting the activation voltage, altering maximal current and/or by slowing inactivation ^43^. Our data shows that deletion of CDKL5 results in loss of Cav2.3 S15 phosphorylation, leading to GoF due to slowed inactivation and amplified gating modulation, similar to the effects observed in Cav2.3 variants. Our results establish Cav2.3 overactivity as a common feature of CACNA1E (DEE69) and CDKL5 (DEE2) epileptic encephalopathies, potentially explaining overlapping clinical features of these diseases.

Cytoplasmic N-terminal S15 phosphorylation of Cav2.3 has been previously detected in global phosphoproteomics screens from brain (phosphosite.org), but the responsible kinase and the role of this phosphorylation were unknown. N-terminal deletions in Cav2.3 ^59^ or the related Cav2.2 ^79, 80^, result in channels with normal gating and conduction properties. Inactivation kinetics were not measured in these studies, yet interestingly Cav2.2 N-terminal deletion mutants seemed to have slower decaying currents, indicating that this region participates in the regulation of inactivation ^79^. Analogously, loss of N-terminal Ser14/15 phosphorylation slows Cav2.3 inactivation in HEK293 cells and neurons, underscoring a previously overlooked regulatory role of the Cav2.3 N-terminus. This domain may interact with Cavβ, as previously suggested for Cav2.2 ^80^.

We also find that cholinergic stimulation amplifies Cav2.3 GoF in absence of CDKL5 phosphorylation, by prominently shifting activation V_1/2_ and increasing currents at maximal activation voltages, thus revealing a second molecular mechanism by which loss of pS14/15 enhances Cav2.3 function. It is known that muscarinic enhancement of Cav2.3 depends on Gαq signalling and PKC phosphorylation at multiple sites, including the intracellular loop I-II ^61^, the most prominent site of α1-β interaction ^53^. We show that loss of pS14 interferes with PKC enhancement upon carbachol application, but the mechanisms of this interplay are yet to be established.

Our recordings from hippocampal neurons in Cav2.3 S15A mice demonstrate that S15 regulates Cav2.3 function in neurons. We show that pS15 is critical for R-type current inactivation and regulation of firing by cholinergic stimulation. Isolating Cav2.3 currents is challenging due to space clamp limitations and the need for inhibitors of other voltage-gated channels. These drugs may have promiscuous effects on Cav2.3 ^81^ or incompletely block other Ca^2+^ currents ^55^. We observed enhanced neuronal excitability and increased pro-epileptiform cholinergic ADPs/DPPs ^58, 63^ in Cav2.3 phosphomutant mice, an effect that was not linked to carbachol suppression of AHPs or other intrinsic excitability properties in these cells. Instead, we suggest that the GoF of unphosphorylated Cav2.3 underlies the hyper-excitability seen in response to muscarinic stimulation. Interestingly, CDKL5 KO mice have altered cholinergic tone ^82^ and the muscarinic antagonist Solifenacin was shown to ameliorate network defects in CDD patient-derived neurons ^83^, supporting an enhanced cholinergic modulation in this human model.

We find striking behavioural similarities between our Cav2.3 phosphomutant mice and CDKL5 KOs. Specifically, S15A mice show reduced encoding and retention of fear memory, a robust phenotype found in full body ^15, 73, 84^ and excitatory neuron-specific ^85^ CDKL5 KO mice. Cav2.3 KO mice display enhanced contextual fear conditioning ^37^. Cav2.3 S15A mice recapitulate the reduced home-cage locomotion behaviour in CDKL5 KO mice ^48, 73^. The deficits observed in voluntary wheel in Cav2.3 phosphomutants could be indicative of mild hypotonia ^86, 87^. Finally, reduced social interaction in Cav2.3 phosphomutant mice was also observed in CDKL5 KO mice ^15, 48, 73^ and inhibitory neuron specific deletion of CDKL5 ^88^. Importantly, these phosphomutant mouse phenotypes mirror human clinical manifestations of CDD, arguing that slowed inactivation and enhanced cholinergic modulation of unphosphorylated Cav2.3 may contribute to these symptoms.

Cav2.3 KO mice exhibit reduced susceptibility to chemical absence seizures ^89–91^ and hippocampal KA-induced seizures ^51^. In agreement with Cav2.3’s role in epilepsy, we find increased seizure susceptibility in females in the low-dose KA injection paradigm in our Cav2.3 S15A mice. The difference between males and females, observed in both ECoG and behavioural seizure analysis, may arise from increased basal function of Cav2.3 in the female brain downstream of hormonal signals. Therefore, gender should be taken into consideration in future mouse CDKL5 KO studies, where seizure susceptibility has not been observed in one gender. In CDKL5 KO males, KA-induced seizure susceptibility changes have not been observed ^18, 48^. Reduced latency to seizure initiation with pentylenetetrazole (PTZ, a GABA_A_ receptor inhibitor) injections has been reported in CDKL5 KO males ^84^ as well as heterozygous females ^74^. Finally, increased severity of seizures upon NMDA injections were observed in male CDKL5 KO mice due to increase NMDAR function ^18^, however NMDAR function is not altered in CDKL5 KO rats ^29^. In absence of pro-convulsants, disturbance-associated seizures are observed only in aged heterozygous females in two different CDD mouse models ^75^, an occurrence that may be linked to developmental changes in Cav2.3 regulation.

In rodents, it is possible to reverse adult phenotypes with reintroduction of CDKL5 genetically ^72^ or using AAV viruses ^92^, indicating an open therapeutic window for treatment. Here, we report a novel functional regulation of Cav2.3 by CDKL5 phosphorylation, which impinges of neuronal excitability. Thus Cav2.3 GoF in absence of CDKL5 may be an important molecular mechanism in CDD pathology. In addition to CDKL5 and CACNA1E encephalopathies, increased Ca^2+^ influx through Cav2.3 may contribute to seizures in Fragile X syndrome, as reduced FMRP function leads to increased Cav2.3 expression ^93^. Cav2.3 also increases susceptibility to drug-induced Parkinson’s disease ^95, 96^. We propose that Cav2.3 inhibitors/ modulators could be beneficial for CDD patients and potentially for a broader population of neurological patients in the future.

## Methods

### Mouse handling and mutant generation

Animals were housed in a controlled environment with 12-hour light/dark cycle. They were fed *ad libitum* and used in accordance with the principles of the “3Rs” and the Animals (Scientific Procedures) Act 1986 of the United Kingdom. Protocols and procedures were approved by The Francis Crick Institute and UCL Institute of Neurology oversight committees. Except for some *in vivo* experiments, mice were housed in groups. Mouse strains were backcrossed into C57 Bl/6 (Jackson) and none of the experimental mice were immunocompromised. Male and female mice aged embryonic day (E)16.5 to postnatal 30 weeks were used as specified.

CDKL5 KO animals were a kind gift from Cornelius Gross ^48^. Mutant Cav2.3 Ser15Ala animals were created in-house using the CRISPR/Cas9 system. The guide contained the sequence 5’AGGCUCAGGCGAUGGAGACU3’. This replaced the original DNA sequence of 5’ AGGCGCAGGCGATGGAGACT 3’ in CACNA1E. Genetically altered C57Bl6 embryos were obtained using electroporation as part of Francis Crick Institute’s Genetic Modification Services. Genotype was determined by in-house PCR or Sanger sequencing (Source Bioscience).

### SILAC phosphoproteomics

#### Sample preparation

Mouse cortical neurons were cultured from individual male E16.5 embryos from a heterozygous Cdkl5 female. Cdkl5 +/Y (WT) or -/Y (KO) genotype of embryos was determined afterwards. Neurons were plated at a density of 8x10^6 cells per 10 cm dish coated with 60 μg/ml poly-D-lysine (Sigma) and 2.5 μg/ml laminin (Sigma). Cells were grown in neurobasal media free of L-Arginine and L-Lysine (Invitrogen) supplied with either Lysine 8 (U-13C6, U-15N2) and Arginine 10 (U-13C6, U-15N4) or Lysine 4 (4, 4, 5, 5-D4) and Arginine 6 (U-13C6). L-Proline (200mg/ml) was supplemented to prevent Arginine to Proline conversion. At DIV4, 1 μM Ara-C was added to inhibit glia growth. For every embryo, one dish was labelled with K8R10 (heavy) and one dish was labelled with K4R6 (light).

At DIV12, SILAC labelled primary neurons from 3 Cdkl5 WT and 3 Cdkl5 KO animals were lysed in lysis buffer containing 20 mM Tris pH7.5, 100 mM NaCl, 10 mM MgCl_2_, 0.25% IGEPAL NP40, 0.5 mM DTT, 1x Protease Inhibitor Cocktail, 1 μM Okadaic Acid. Each sample was checked for SILAC label incorporation and Arginine to Proline conversion. Lysates were mixed to obtain 3x heavy WT/light KO and 3x heavy KO/light WT samples containing 1 mg total protein. Proteins were digested overnight with Sequencing Grade Modified Trypsin/Lys-C (Promega) at 37^0^C, desalted using Sep-Pak Classic C18 Cartridges (Waters) and vacuum dried completely. Samples were enriched for phosphopeptides using Titansphere titanium dioxide beads (GL Science) in batch mode with the following buffers: Loading buffer: 80% ACN, 5% TFA, 1M Glycolic Acid, Wash buffer 1: 80% ACN 1% TFA, Wash buffer 2: 10% ACN, 0.2% TFA, Elution buffer 1: 1% Ammonium Hydroxide, Elution buffer 2: 5% Ammonium Hydroxide. Phosphopeptides were cleaned up using C18 Stage Tips with a centrifuge protocol and vacuum dried completely. Samples were stored at -80^0^C until required for analysis by mass spectrometry.

#### Mass spectrometry

Each sample was resuspended in 35 μl 1% TFA, sonicated for 15 minutes and injected 3 times (10 μl per injection). Peptide mixtures were separated on a 50 cm, 75 μm I.D. Pepmap column over a 3 hr gradient and eluted directly into the mass spectrometer (Orbitrap Velos). Xcalibur software was used to control data acquisition. The instrument ran in data dependent acquisition mode with the top 10 most abundant peptides selected for MS/MS by CID, MSA or HCD fragmentation techniques (one fragmentation technique per replicate).

Data processing was performed using the MaxQuant bioinformatics suite (v 1.3.0.5) and protein database searching was done by the Andromeda search engine using a Uniprot database of *Mus musculus* proteins amended with common contaminants. Default search settings were used including an FDR of 1% on the phosphosite, peptide and protein level and matching between runs for peptide identification. K4R6 and K8R10 were set as labels with a maximum of 3 labeled AAs. Phosphorylation at serine, threonine or tyrosine residues, oxidation (Met) and protein N-terminal acetylation were set as variable modifications. Carbamidomethylation of Cys-residues was set as a fixed modification.

Data was further processed in Perseus v.1.6.15.0. Phospho (STY) sites from the six experimental samples in the MaxQuant search were loaded and site tables expanded. Potential contaminants and reversed hits were removed. Normalized H/L ratios were log2 transformed and filtered for rows containing at least three valid values. Potential CDKL5 substrates were annotated using the R-P-X-pS/pT motif ^20^. A one-sample t-test was performed comparing the normalized ratios to the null hypothesis and visualized as a volcano plot.

### Mutagenesis

Point mutations were introduced using QuickChange site-directed mutagenesis (Agilent). Complimentary primers contained the desired mutations flanked by at least 18 bp. To ensure mutagenesis efficiency for each construct, two separate PCR reactions with each primer were set up: 0.2 μM primer, 100 ng template DNA, 0.2 mM dNTP mix, 1x Pfu buffer and 2.5 U Pfu Ultra HF. The PCR programme was run 4x, after which complementary samples were combined and the PCR run repeated 20x. PCR products were treated with DpnI for 1hr at 37^0^C and transformed into XL-10 Gold competent cells. At least 4 colonies were used for each DNA miniprep (QIAprep, Qiagen) and mutations were confirmed by Sanger sequencing (Source Bioscience). The constructs used were as follow: 1) HA-tagged human α1E subunit (gene bank L27745.2, gift from L. Parent, Université de Montréal), in a commercial vector under control of CMV promoter ^97^. 2) The N-terminal FLAG-tagged full-length human CDKL5_107_ and N-terminal HA-tagged human CDKL5_1-352_ kinase domain construct kinase as previously described ^20^.

### Cell line maintenance and transfection

Tetracycline-inducible HEK293 cells expressing human Cavβ3 and α2δ1 subunits were obtained from SB Drug Discovery (Glasgow, UK). HEK293 cells stably expressing human Cav β1b and α2δ1 subunits were kindly provided by Andrew Powell (GSK Ltd., Middlesex, UK). Naïve HEK293T cells were supplied by Francis Crick Institute’s Cell Services.

All cells were cultured in high glucose DMEM with 10% Fetal Bovine Serum (tetracycline free, Clontech or Gibco) and penicillin/streptomycin 50units/ml-50 mg/ml. They were passaged using TrypLE Express and maintained under selection antibiotics as appropriate: HEKβ3/ α2δ1, Zeocin 300mg/ml (Invivogen) & Blasticidin 5mg/ml (Invivogen); HEKβ1b/α2δ1, Puromycin 1mg/ml and Hygromycin-B 200mg/ml. Cell culture reagents were obtained from Gibco/Invitrogen unless specified.

Selection antibiotics were removed before transfection and doxycycline 1μg/μl added in the case of β3-expressing cells. Transient transfection was carried out using X-tremeGENE^TM^ 9 reagent (Roche) following manufacturer’s instructions. Total DNA was 1.5-1.6μg. For Western blot experiments, human α1E-HA subunit and CDKL5 (HA-kinase domain or FLAG-full length) were co-transfected in a 1:1 ratio, substituting kinase for empty pcdna3 vector in some cases. For electrophysiological recordings α1E-HA, FLAG-CDKL5 full length and GFP (pcdna3) were co-transfected at a ratio of 10:4:1 (HEK β3 cell line) or 8:4:1 (HEKβ1b cell line). To study muscarinic regulation, human α1E-HA, FLAG-CDKL5, muscarinic receptor type 3/1 (CHRM3/1, Addgene) and GFP were co-expressed at a ratio of 8:4:3:1. For dopamine modulation experiments, human α1E-HA, FLAG-CDKL5 and dopamine D2 receptor (gfp-DRD2, Addgene) were co-transfected at a ratio of 2:0.75:1.

### Western Blotting

Three different types of sample were used for Western blot experiments: 1) HEK293 cells 48 hr post-transfection, 2) Mouse brain tissue collected after cervical dislocation, followed by dissection and snap-freezing in liquid nitrogen, 3) Human iPSC derived neurons obtained as frozen pellets from Cleber Trujillo (UCSD, USA). Protocols for iPSC generation, patient *CDKL5* mutations and related control details are described in ^83^. All lysates were prepared in 1x/2x sample buffer (Invitrogen) with 0.1-0.2 M dithiothreitol (Sigma). This was followed by sonication, centrifugation at 13000g and denaturation at 70^0^C, 10 min. Protein was loaded onto NuPAGE 8% Bis-Tris polyacrylamide gels (Invitrogen) for electrophoresis and subsequently transferred to polyvinylidene difluoride membranes (Millipore) for 15-22 hr at 20V in 10% MetOH, Tris Glycine buffer. Membranes were blocked for 30 min in 5-10% skimmed milk and incubated with primary antibodies overnight at 4^0^C or 1hr at room temperature (RT). Secondary incubation in horseradish peroxidase-conjugated (HRP) antibodies was for 1hr at RT. Chemiluminescence signals were detected using Amersham ECL (Cytiva) and Amersham Imager (GE Healthcare). Analysis was perfomed using Fiji 2.1. Phospho-S14/15 channel levels were measured as the ratio between pS14/15 Cav2.3 signal and total Cav2.3 signal. Total channel levels are expressed as the ratio over loading control GAPDH. Each replicate in a blot constitutes a data point normalized to the average internal control signal in each blot. At least two independent immunoblot experiments with technical replicates were used for quantification, as specified in figure legends.

Primary antibodies used in each sample were: mouse anti-HA 1:2000 (Biolegend, 901513; HEK293), rabbit anti Cav2.3 N-terminus 1:250 (Covalab, custom 1, HEK293 & brain), rabbit anti pS15 Cav2.3 1:500 (Covalab, custom 2, HEK293, brain & human neurons), mouse anti Cav α1E 1:1000 (Synaptic Systems, 152411, human neurons), rabbit anti CDKL5 1:2000 (Atlas, HPA002847, HEK293), mouse anti GAPDH 1:50000 (Abcam, ab8245, brain & human neurons). Custom made Cav2.3 Ser15 phosphospecific antibodies were raised by immunizing rabbits with peptide PRPG(pS)GDGDSDQSRNC. Phosphorylated Cav2.3 antibody was obtained after double purification. First, they were selected against non-phosphorylated peptide PRPGSGDGDSDQSRNC. This antibody is used as total Cav2.3 antibody (custom 1). Next the eluted fraction was purified again by binding to PRPG(pS)GDGDSDQSRNC -linked beads, obtaining pSer15 Cav2.3 (custom 2). Secondary antibodies used were: donkey anti rabbit HRP 1:10000 (Jackson, 711-035-152), donkey anti mouse HRP 1:10 000 (Jackson, 715-035-151).

### *In vitro* electrophysiology

#### HEK293 cells

Cells were split 24hr after transfection and replated on glass coverslips and allowed to recover overnight. Whole-cell patch clamp recordings in isolated cells were performed 48-72 hr post-transfection at room temperature. Transfected cells were identified by GFP fluorescence. Data were sampled at 20kHz and filtered at 1-2 kHz. Cells were continuously perfused (1ml/min) with extracellular solution containing (in mM): NaCl 120, TEACl 20, HEPES 10, glucose 10, KCl 5, MgCl_2_ 1, CaCl_2_ or BaCl_2_ 5, pH 7.4 with NaOH. Pipettes (2.5-3.5 ΩM) were pulled from borosilicate glass (1.2x0.69mm, Harvard Bioscience) and filled with the following intracellular solutions adapted from ^40^ to minimize current run-down (in mM): CsMeS 140, EGTA 5, MgCl_2_ 0.5, MgATP 5, HEPES 10, for GPCR experiments; CsMethanesulfonate 125, TEACl 5, EGTA 5, MgATP 5, Na_3_GTP 0.3, Na-phosphocreatine 5, Na-pyruvate 2.5, HEPES 10, pH 7.25, 285-295 mOsm, all other recordings. Upon seal break, currents were evoked every 5/10s with a step from -80/-100mV (holding potential, HP) to 0/+10mV. After an initial run-up period, steady state current voltage (IV) relationships were obtained with 100ms depolarising test steps to +60/+80 in 10mV increments every 10s. For voltage dependence of inactivation the protocol consisted of a 1s pulse from -140 to +20mV in 10 mV increments, followed by a 50 ms test pulse to 0 mV, every 10s. Peak currents (I_peak_) in response to test voltages (V_test_) were used to plot IVs and measure the voltage dependence of activation and inactivation. Reversal voltage (E_rev_) was estimated by extrapolating a linear fit to the IV curve from +20-+40 mV. Conductance (g) was calculated using the equation (g=(I_peak_)/(V_test_-E_rev_)). Voltage of half-maximal activation or inactivation (V_1/2_) were measured by fitting with a Boltzmann equation (Y= A+(B-A)/(1+exp((V-V_1/2_)/K)) where Y is the conductance or the current, and A and B are the minimum and maximum amplitudes of the fit. Time course of open state inactivation (t_inact_) was estimated by fitting a single exponential function to current decay in response to test pulses. The average of 3 fits at each voltage was used. For all transfection conditions, cells with outward currents >40% of the peak current at +10mV or with τ_inact_ that did not decay uniformly with depolarization were excluded from analysis. Similarly, cells were not used for current density or τ_inact_ comparisons if run-down was >30% of the maximal current recorded in the experiment as this is known to affect inactivation time ^40^. To examine recovery from inactivation, an inactivating pulse to +10mV of 0.8-1s (pulse 1), was followed by a variable recovery period (Δ150-200ms) and a test pulse to +10mV (pulse 2), at 0.1-0.3 Hz. Time course of recovery was calculated using the ratio of I_peak_ in response to pulse 2/pulse 1 plotted against duration of recovery period and fitted with single exponential equation. Carbachol, quinpirole and PMA were bath applied at ∼3 ml/min whilst activating Cav2.3 currents with brief monitoring (25ms) test steps to 0 or +10mV, every 2, 5 or 10s, depending on speed of run-down. Upon steady state, IVs were obtained. Cells included in the analysis showed no significant changes in series resistance and a consistent shift in activation V_1/2_, with current amplitude matching between IV and brief monitoring steps. Leak and capacitance subtraction was applied to all recordings using -P/5 protocol and series resistance compensated from 75-95% to keep voltage error <5mV.

### *Ex vivo* electrophysiology

#### Hippocampal slices

Mice older than P10 were anaesthetized by intraperitoneal injection of ketamine (80mg/kg) and xylazine (10mg/kg). To avoid bias, the genotype was unknown at the time of experiment. Animals were decapitated and the brain transferred to ice cold artificial cerebrospinal fluid (standard aCSF) containing (in mM): NaCl 125, NaH_2_PO_4_ 1.25, NaHCO_3_ 26, KCl 2.5, glucose 25, MgCl_2_ 1, CaCl_2_ 1, saturated with carbogen. Cerebellum and olfactory bulb were discarded. Coronal or transverse hippocampal slices (300μm) were prepared with a Leica VTS 1200S vibratome and transferred to an immersion storage chamber containing bubbled aCSF at 35^0^C for 30 min. Slices were allowed to recover for a further 30 min at room temperature (RT) before electrophysiology experiments. Somatic whole-cell patch clamp recordings were obtained from CA1 pyramidal neurons using borosilicate electrodes (3-4.5 MΩ). Series resistance (R_s_) and input resistance (R_in_) were monitored throughout recordings using a 100ms, -5mV step from -60mV to evoke passive membrane responses. R_s_ was calculated from the amplitude of the capacitive transient and R_in_ from the steady state currents. These were sampled at 20kHz and filtered at 10kHz. The maximum change in Rs permitted was 25%.

*R-type Ca^2+^ currents* - Recordings were obtained from young neurons (P10-11) no longer than 6h post slicing using the blind technique. Currents were recorded at RT from P10-11 coronal slices during bath application of a modified aCSF solution (in mM): NaCl 115, TEACl 10, NaHCO_3_ 26, KCl 2.5, NaH_2_PO_4_ 1.25, glucose 25, MgCl_2_ 1, CaCl_2_ 2; and in presence of the following inhibitors (in μM): TTX 0.5, gabazine 1, APV 25, NBQX 5, 4AP 5, Nifedipine 10 (Tocris), w-conotoxin GVIA 2, w-agatoxin IVA 0.2, w-conotoxin MVIIC 2 (Alomone) and BSA 1mg/ml. Drug stock solutions were prepared in water and stored at -20^0^C. External solution was constantly bubbled and re-circulated. Slices were exposed to ion channel inhibitors for at least 20 min before voltage clamp acquisition. Patch pipettes were filled with (in mM): CsMethanesulfonate 130, TEACl 10, Na-phosphocreatine 5, MgATP_2_ 4, Na_3_GTP 0.3, HEPES 10, EGTA 2 and biocytin 0.2% pH 7.3 with CsOH. After 10 min stabilization in the whole-cell configuration, IV curves were obtained with voltage steps from -80 to +40, HP=-70mV. A -P/5 leak and capacitance subtraction protocol was used and R_s_ was compensated 65-85%, to minimize voltage error. Capacitance (C_m_) was calculated using equation: C_m_ [pF] = τ_m_ [ms]/R_s_[MΩ], where τ_m_ is the decay time constant of capacitative transients. Time course of open state inactivation was estimated by iterative single exponential fit to current decay. A positive correlation was found between τ_inact_ and C_m_ for C_m_<80pF, thus this was established as the cut off cell size criteria for inclusion in τ_inact_ comparisons. Cells with large cells outward currents at depolarising voltages (>50% of the peak current at +10mV) were also excluded. *Small-conductance Ca^2+^-activated (SK) currents and excitatory postsynaptic currents (EPSC)*-Recordings were obtained from adult neurons (5-6 weeks) in transverse slices. SK currents were measured as tail currents following a 100ms step from -50 mV to +10mV (0.3 Hz) to engage VGCCs. Standard aCSF (CaCl_2_ 2mM, RT) was supplemented with (in μM): TTX 0.5, TEA 1000 and XE991 5, to isolate SK currents by inhibiting temporally overlapping voltage-gated Na^+^ and K^+^ conductances. The intracellular solution was (in mM): KGluconate 135, KCl 10, Na-phosphocreatine 10, MgATP_2_ 2, Na_3_GTP 0.3, HEPES 10, pH 7.3, supplemented with 8-cloro-phenylthio cAMP 50μM, to inhibit the slow afterhyperpolarizing current. SK current identity was confirmed at the end of all experiments by reversible d-tubocurarine (dTC, 50μM) inhibition. dTC-subtracted currents were used for quantification. Traces were sampled at 5 kHz and filtered at 1kHz. SK current amplitude was measured at the peak of the after-current and charge as the integral 500ms post stimulus. EPSCs were isolated at -70 mV in standard aCSF (1) (CaCl_2_ 2.5mM, 30-32^0^C) with gabazine 1 μM and Kgluconate-based internal solution. Traces were sampled at 10kHz and filtered at 2kHz. Upon 5 min stabilization in whole-cell, a 2 min gap free trace was analysed using template-based event detection and threshold match >3.

*Current clamp recordings* - Recordings were obtained from adult neurons (5-6 weeks) in coronal slices in standard aCSF (2mM Ca^2+^, 30-32^0^C) and Kgluconate-based internal solution. Membrane potential (V_rest_) was measured throughout the recording with 0 current injection and maintained around -65mV during stimulation. Input-output relationships were obtained with 1s current injections from -100 to 450 pA in 50 pA increments. Traces were sampled at 10kHz and filtered at 5kHz. Action potentials were identified with threshold-based event detection (0mV) and amplitude measured from V_rest_. Spike attenuation during the train was calculated as ratio between last and first spike amplitudes. Depolarizing plateau potentials (DPPs) were defined as sustained depolarizations >20mV upon stimulus termination. Medium duration afterdepolarizations (ADPs) were measured 100ms post-1s step depolarization. Firing patterns were compared using binomial tests or comparisons of non-linear least squares regression fits to the data distribution. The effect of carbachol on the medium and slow afterhyperpolarizations (mAHP and sAHP) was used as positive control for muscarinic receptor activation. AHPs were evoked at -65mV by a burst of five 2nA, 2ms somatic current injections at 100Hz every 20s and measured at the peak and 500ms post stimulus respectively.

All patch clamp data was obtained with Multiclamp 700B, Digidata 1440A and pClamp^TM^ 10 software (Molecular Devices). Data were analysed using Clampfit 10.7/11.2 and OriginPro 9.8. Liquid junction potential was not corrected.

### *In vivo* electrophysiology

#### Chronic electrocorticogram recordings

Surgical procedures were performed in adult male and female mice (8-10 weeks old) using a stereotaxic frame (Kopf) under isoflurane anaesthesia and temperature control. Each animal was subcutaneously implanted with an electrocorticogram (ECoG) transmitter (A3022B-CC-B45-B, Open Source Instruments, Inc.). The recording electrode was placed above the visual cortex (AP -2, ML 1.5) and the ground electrode in the contralateral prefrontal hemisphere (AP 1.8, ML 1.5). Implanted WT and Hom S15A animals (total 18) were housed individually and no drug treatment was given. The ECoG was recorded wirelessly (sampling frequency 512 Hz, band-pass filter 1-160 Hz) for two weeks using software from Open Source Instruments, Inc. Simultaneous video recordings were obtained (6x/hour) using IP cameras from Microseven (https://www.microseven.com/index.html). The coastline, kurtosis and power spectrum analysis (1-160Hz) were performed using Python. Researchers were blinded to animal genotype during data acquisition and analysis.

#### Kainic acid susceptibility

At the end of the chronic recordings, low dose 5mg/kg kainic acid (Tocris Bioscience) dissolved in sterile saline was administrated intraperitoneally every 30 minutes to assess brain susceptibility to increasing dosage of chemo convulsant. The clock started at the first injection from 0 minute. Seizure severity was initially assessed while the experiment was ongoing at 10-minute intervals, using a modified Racine scale: 1. immobility; 2, hunched position with facial jerks, 3. rearing and forelimb clonus, 4. persistent rearing and forelimb clonus, falling, 5. generalized tonic clonic convulsions, or wild jumping ^78^. These assessments were confirmed by re-analyzing the video recordings. Latency measures the length of time from the start of the experiment to the first indication of generalized seizure (Stage 5 on Racine scale). Both kainic acid inductions and analysis were performed by a researcher blinded to the genotypes. A total of 17 implanted mice were used for susceptibility experiments, one male mouse was removed from the study due to surgical complications with the transmitter. In a subset of these where transmitter performance was not compromised, ECoG recordings during seizure inductions were recovered and used for quantification. Traces were analysed using semi-automated seizure detection as described previously ^77^. The cumulative coastline was calculated from when the first epileptiform activity was detected until the mice reached generalized seizure (stage 5).

### Behavioural analysis

#### Home Cage monitoring

Two separate cohorts of WT and homozygous S15A Cav2.3 mice of both sexes and aged between 12-16 weeks were used. Animals were housed individually and monitored using the Digital Ventilated Cage system (DVC^®^,Techniplast, Italy). The DVC consists of standard size, barcoded IVC units equipped with a 3x4 arrangement of cage-floor electrodes. Capacitance sensing-based detection of animal position allows for real-time tracking of locomotion activity 24/7. Following a 2-week-acclimatization period, baseline activity was recorded and analysed. Rotating wheels (Ø 4.4 in) were then introduced, and animals allowed to familiarize themselves for another week before data collection. All activity metrics were calculated using the DVC Analytics 3.4.0 platform ^98^. For locomotion index, the average signal of all 12 cage electrodes was used. Data were grouped in 60 min bins for each day.

#### Behaviour tests

Behaviour experiments were conducted in 8-week-old WT and homozygous S15A Cav2.3 littermates of both sexes, housed under an inverted light cycle. Mice were handled daily for one week prior to experiments. All tests were video recorded for off-line analysis. Tracking data were analysed using Ethovision XT 15 (Noldus Information Technology, Netherlands) equipped with three-point (nose, body centre, and tail) detection settings. One cohort of mice was used for all 7 tests, carried out in the order described below. Mice were allowed to habituate to the testing room for at least an hour before the behaviour test.

Hind-Limb clasping – Mice were suspended by the middle of the tail and lifted 15 cm above the ground; the extent of hind-limb clasping was recorded for 15 seconds. The room lighting was kept at 90 Lux. Hind-limb clasping was scored from 0 to 3: score 0 = both hind limbs were splayed outward away from the abdomen; score 1 = one hind limb retracted inward toward the abdomen for at least 50% of the observation period; score 2 = both hind limbs partially retracted inward toward the abdomen for at least 50% of the observation period; score 3 = both hind limbs completely retracted inward toward the abdomen for at least 50% of the observation period. Hind-limb extension reflex severity scores were calculated by averaging three trials separated by 1 min.

Open field test - The apparatus consisted of an illuminated (70 lux) white floor PVC foam arena (50 x 50 x 40 cm). A central zone (16 x 16 cm) was pre-defined. Test mice were placed in the centre of the open field and allowed to explore the arena for 30 minutes. At the end of each trial, the mouse was returned to the home cage and the arena was cleaned. Total distance travelled in 30 minutes and percentage of time spent in the central zone were used as measures of locomotor activity and anxiety levels, respectively.

Accelerating Rotarod test - Day 1: Mice walked on a rotating rod (Ugo Basile model 47650, Italy) at constant speed (4 rpm) for three minutes for two acclimatization trials. Day 2: Each mouse was placed on the rotating rod for three test trials, during which the rotation speed gradually increased from 4 to 40 rpm within four minutes. The inter trial interval was 1 hr. Performance was evaluated by measuring the latency to fall.

Three-chamber social interaction test – The apparatus consisted of a white rectangular PVC foam arena (62x42x23 cm) divided into one central and two adjacent compartments of equal size compartments connected by square openings. Empty cylindrical wire cages (Ø 8 cm, height 18 cm) were placed in the lateral chambers. Each mouse was placed in the central chamber and allowed to explore the arena for 10 min. The brightness of the room was kept at 10 lux. Chamber occupancy during habituation was equal between groups. After this, the mouse was returned to the waiting cage for 3 minutes. To assess social preference, an unfamiliar WT mouse of the same age, sex and strain (stranger 1) was gently introduced inside one of the wire cages to serve as a social stimulus. An unfamiliar and inanimate object was added to the other wire cage as the non-social stimulus. The test mouse was placed in the apparatus containing the social and non-social stimuli for 10 minutes. After that, the test mouse was removed and kept in the waiting cage for 3 minutes. All compartments were cleaned with 30% ethanol solution before the next test phase. To assess preference for social novelty, the non-social object from the previous test phase was replaced by an unfamiliar mouse (stranger 2) and the previous stranger 1 (now familiar mouse) was placed into the wire cage of the opposite compartment. The test mouse was placed in the central chamber and allowed to explore the arena for the next 10 minutes, after which it was returned to the home cage. The location of the novel and familiar mice with respect to the side compartments was counterbalanced across trials. The exploration of the target mouse/object was scored when the mouse’s nose was detected within 2 cm from the cylindrical wire cage.

Y-maze spontaneous alternation test – Testing occurs in a Y-shaped maze with three arms (1,2,3) made of opaque plastic (34 x 9 x 14 cm) oriented in a 120^0^ angle. Each mouse was randomly placed into one arm and allowed to explore the maze for 10 minutes. Room lighting was kept at 10 lux. An arm entry was recorded manually when the mouse moved beyond the central triangle of the maze and entered an arm with all four paws. Alternation behaviour was defined as consecutive entries into each of the three arms in overlapping triplet sets (e.g.: 1, 2, 3 or 2, 1, 3 or 3, 2, 1). The percentage of alternations was calculated as the number of actual alternations divided by the maximum possible number of arm entries.

Barnes-maze test – Twenty-four hours before training, the animal was habituated to the apparatus and escape box for a minute. During the training phase (4 days), mice were trained to locate the target hole (with an escape chamber underneath) from among 20 holes evenly spaced around the perimeter of an elevated circular open field (Ø 96 cm). Each animal was initially placed in the centre of the arena covered by an opaque cylinder, which was removed 10-20s after the start of the trial. Room lighting was kept at 635 lux. The mouse is then free to explore the platform for 3 minutes using four visual cues to aid navigation. If the mouse does not enter the escape chamber within 3 minutes the experimenter guides the mouse gently to the escape box (20x11x7cm) and leaves the mouse inside for 1 min. This step is repeated two more times a day (3 trials in total per day) with at least 20 minutes in between where the mouse is placed back in its home cage. To assess learning, a probe test in carried out 24h after the last training, where the escape box has been removed from the target hole. The mouse is placed on the platform and free to explore it for 3 minutes. Distance to first: distance walked from the centre of the arena to the target hole (centre-point detection); latency to first: time to reach the target hole from the centre of the arena (centre-point detection); errors to first are defined as checking any hole before reaching the escape box (nose-point detection).

Contextual fear conditioning – The fear conditioning chamber (27x27x35 cm, Ugo Basile, Italy) was equipped with a shock grid floor and a digital Near Infrared Red Video Fear Conditioning system. Mice from each genotype and sex were examined in four successive phases comprising: conditioned acquisition (day 1), memory of the conditioned background context A (day 2), exploration in new background context B (day 3) and memory extinction in the conditioned context A (day 4-7).

Context A was characterized by a cubic shape, illuminated at 100 Lux and the presence of a vanilla odour. The floor consisted of metal rods, mediating the foot shock. Context B was also cubic shaped, not illuminated and without additional odour cues. On day 1: Mice were introduced in the conditioning chambers scented with vanilla odour for 8 min and received 3 unconditioned stimuli (US: 1s, 0.25 mA foot shock, 2 min inter-trials interval). To examine the conditioned response to the context, mice were reintroduced in the training context 24 h later (day 2) and monitored for 5 minutes in the absence of the US. On day 3, the mice were introduced to a new context (context B) for 5 min to examine whether freezing behaviour is associated to a specific context (context A). During days 4-7 mice were placed back in context A without US to assess extinction of the association of the specific context A and the US (which is described as a form of new learning).

### Statistical Analysis

Statistical analyses were performed in GraphPad Prism 9. Two-tailed Student’s T tests and one-or two-way ANOVA with Geisser-Greenhouse correction followed by Fisher’s LSD test were used for most statistical comparisons, unless otherwise specified. Repeated measures ANOVA was used as appropriate if there were no missing values. Data are presented as mean ± S.E.M. Box and whiskers (min to max) or violin plots are used to illustrate data spread and frequency distribution respectively; percentile 25/75, median and mean are indicated by lines/markers.

### Data availability

The datasets generated and analysed during this study are available from the corresponding authors on reasonable request.

### List of supplementary materials

Supplementary Figures 1-4.

## Acknowledgements

We thank Ultanir lab members for valuable discussion and thoughtful comments. We thank Helen Flynn for valuable assistance with mass spectrometry. We thank Crick GEMS team for creation of CACNA1E S15A mice. This work was supported by the Francis Crick Institute which receives its core funding from Cancer Research UK (CC2037), the UK Medical Research Council (CC2037), and the Wellcome Trust (CC2037); Loulou Foundation Project Grant (11015); Crick-MSD Framework collaboration grant (11202). For the purpose of Open Access, the author has applied a CC BY public copyright licence to any Author Accepted Manuscript version arising from this submission.

## Author contributions

M.S.C., L.L.B., G.L. and S.K.U. designed the experiments. M.S.C. performed and analysed electrophysiology and biochemistry experiments. L.L.B. performed mass spectrometry. Y.Q. and G.L. conducted/ analysed EEG and kainic acid-induced seizure experiments. A.T.L. and M.S.C. performed behaviour experiments/ analysis. L.S., S.M., S.C. provided technical assistance in molecular biology and electrophysiology. M.S.C. and S.K.U. wrote the initial version of the manuscript. J.R. contributed to the original draft and review/editing along with all authors.

## Materials and Correspondence

Correspondence and material requests should be addressed to Sila K. Ultanir and Marisol Sampedro-Castañeda

---

**Supplementary Fig. 1:**
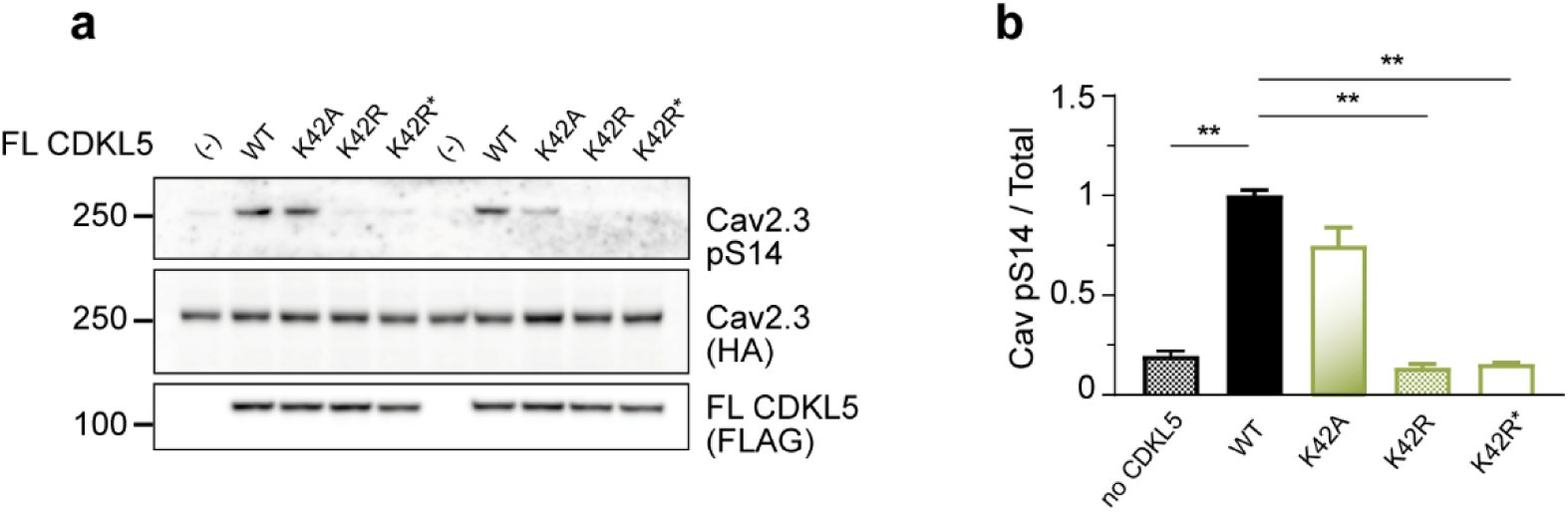
Full-length CDKL5 phosphorylates Cav2.3. **a** Western blot of HEK293 cells stably expressing human β3/α2δ1 subunits and transiently co-transfected with human WT HA-α1E (Cav2.3) and FLAG-CDKL5 full length (FL CDKL5: WT, K42A, K42R, K42R*+DFG-AFG kinase dead mutants). (-) transfection condition excludes CDKL5. The K42A mutation in FL-CDKL5 retains some kinase activity on α1E, while K42R and K42R* mutants are fully inactive and were thus selected for electrophysiology experiments. **b** Quantification of relative phospho-α1E (pS14 Cav2.3) levels for the experiment in **a** (**p<0.0001, Brown-Forsythe & Welch ANOVA, Dunnet’s test; n=4-6, 3 blots, 2 transfections).

**Supplementary Fig. 2:**
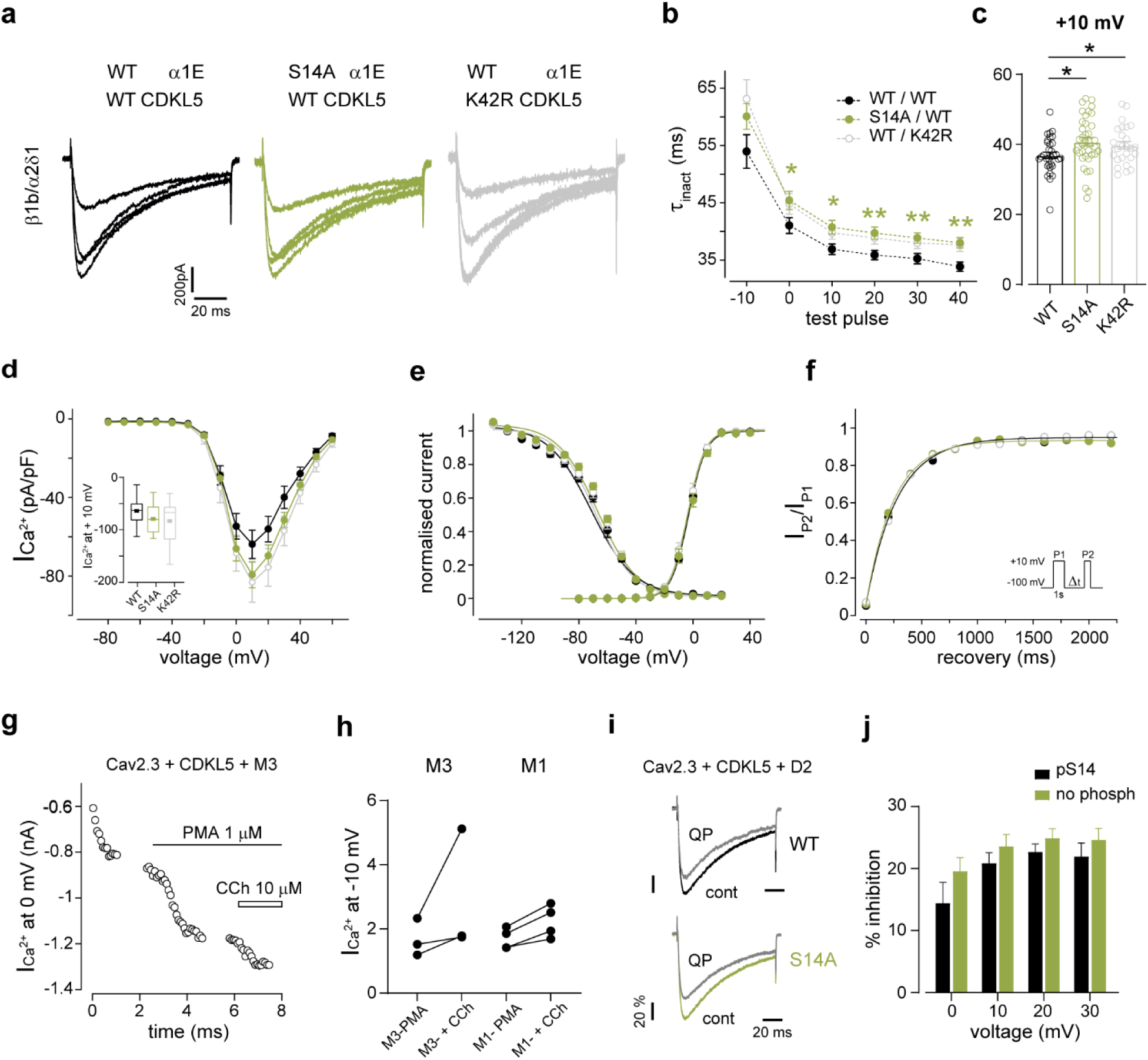
Functional characterization of phospho-Ser14 Cav2.3 in HEK 293 cells expressing β1 accessory subunit. **a** Depolarization-evoked current responses in HEK293 cells stably expressing human β1/α2δ1 subunits and co-transfected with human Cav2.3 (WT α1E or S14A α1E mutant) and FLAG-CDKL5 full length (WT CDKL5 or K42R*+DFG-AFG kinase dead CDKL5 mutant). Colours denote different construct combinations. Traces show steps from -10 to +20mV from -100mV; Ca^2+^ was charge carrier. **b** Open channel inactivation tau (τ_inact_) for Cav2.3 with (WT/WT, n=28-32) and without (S14A/WT, n=34-37; WT/K45R, n=25-26) CDKL5 phosphorylation. WT/WT vs. S14A/WT: *p<0.05, **p<0.01 as indicated; WT/WT vs. WT/K42R*, p<0.01 at +40mV, p<0.05 at - 10,10,20,30mV, Two-Way ANOVA, Fisher’s LSD. Some of the cells included in this plot co-expressed M3 receptors; this was *per se* not found to affect inactivation kinetics. **c** Individual data points for τ_inact_ at +10mV (WT vs. S14A *p=0.014, WT vs. K42R *p=0.0468 two-tailed unpaired t tests). **d** Current-voltage relationship for experiments in **a** (WT/WT, n=15; S14A/WT n=15-17; WT/K42R* n=12-15) and current density at +10mV (inset, p>0.05, OneWay ANOVA). Data was acquired with 100ms voltage steps from -100mV in +10mV increments every 10s. **e** Normalised Cav2.3 conductance and voltage dependence of inactivation for the same three transfection conditions. Activation V_1/2_, n: WT/WT -3±1 mV, 14; S14A/WT -3±1 mV,17; WT/K42R* -2±1 mV, 14; Inactivation V_1/2_, n: WT/WT -66±2 mV,14; S14A/WT -63±1 mV,14; WT/K42R* -65±2 mV,11 (p>0.05, One-Way ANOVA). For inactivation protocol see Methods. Solid lines are Boltzman fits to the average data. **f** Inactivation recovery time using a double pulse protocol with variable inter-pulse recovery time (inset). Recovery tau, n: WT/WT 265±21 ms, 4; S14A/WT 263±21 ms, 5; WT/K42R* 290±15 ms, 4, p>0.05 Kruskall-Wallis, Dunn’s test). **g** Time course of Cav2.3 current amplitude at 0mV in the β1/α2δ1-stable cell line co-transfected with α1E WT, CDKL5 and muscarinic receptor type 3 (M3). PKC activation with PMA did not occlude CCh enhancing effect. **h** Relative change in current amplitude for same experiment as **g** with either M1 or M3 receptors. No muscarinic inhibition was observed. **i** Cav2.3 current at +10mV in β1/α2δ1 cell line co-transfected with α1E (WT or S14A), CDKL5 (WT) and dopamine receptor type 2 (D2). Quinpirole 100nM (QP) application resulted in reliable and reversible inhibition of WT and S14A currents. **j** Quinpirole inhibition of Cav2.3 with (n=7) or without pS14 (n=12-13) (p>0.05 Two-Way ANOVA, Fisher’s LSD). Data for α1E S14A phosphomutant and K42R* CDKL5 conditions (no phosphorylation) were pooled together.

**Supplementary Fig. 3:**
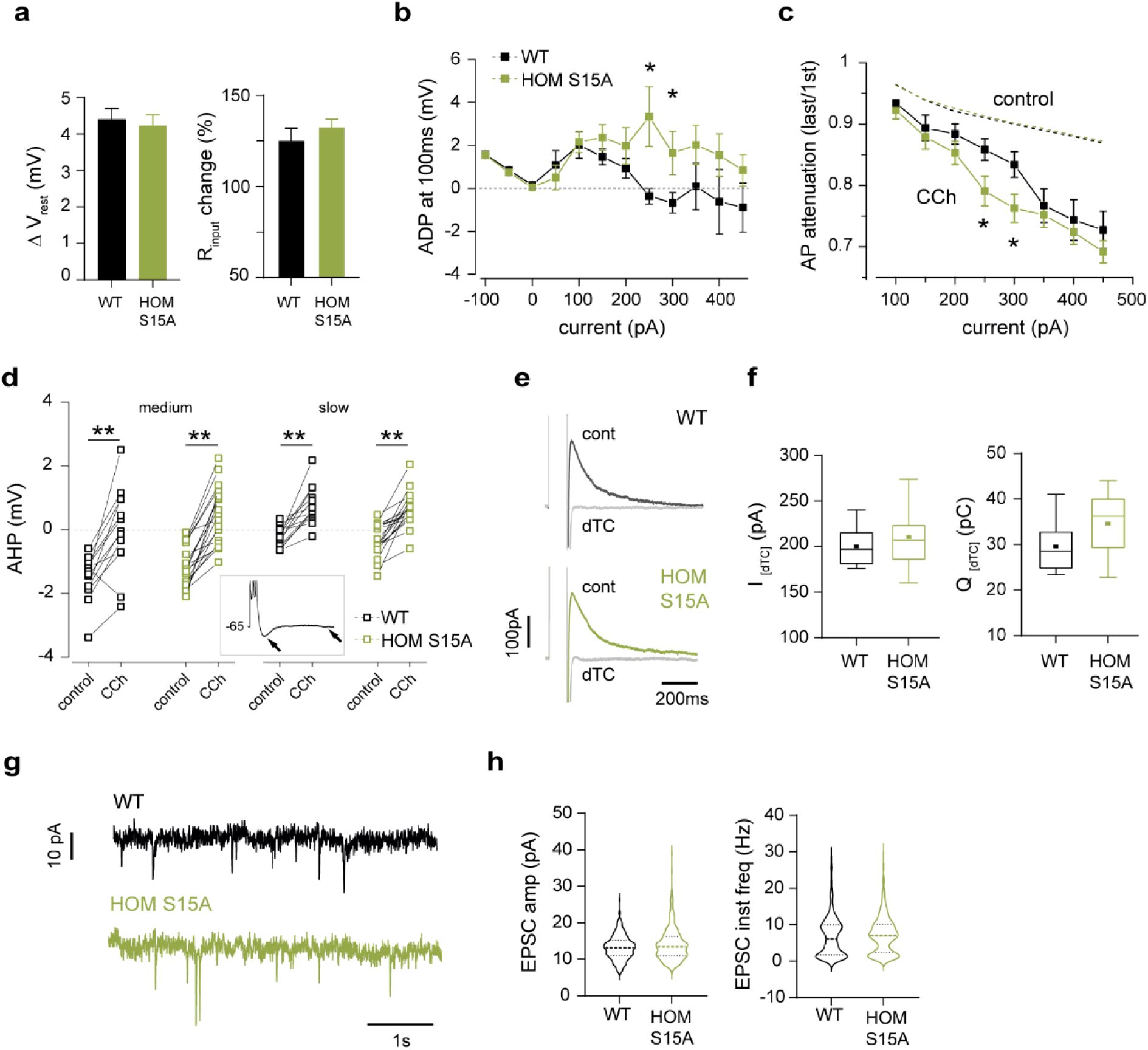
Additional CA1 neuron responses to carbachol and other intrinsic properties in adult WT and HOM S15A mice. **a** Relative changes in membrane potential (V_rest_) and resistance (R_input_) upon 10 μM CCh application in WT (13) and HOM S15A (17) neurons (7 mice/genotype, p>0.05, unpaired t tests). **b** CCh-evoked medium duration afterdepolarization (ADP) measured 100ms post 1s-long stimulus (WT n=13, HOM n=21; p=0.07 Two-way ANOVA; *p<0.05 Fisher’s LSD). **c** Spike attenuation in control (dotted lines) and upon CCh-induced depolarization (markers) shown as ratio between last and first spike amplitude in 1s train (WT n=13, HOM n=19, Two-way ANOVA; *p<0.05 Fisher’s LSD). **d** Medium and slow duration afterhyperpolarizations (AHPs) quantified at the peak and 500ms post stimulus, respectively (inset). AHPs evoked at -65mV by a 100Hz burst of somatic current injections were equal in WT (13) and HOM S15A (18-19) mice (mAHP, -1.5 *vs* -1.2mV; sAHP, -0.2 *vs* -0.3mV, p>0.05, unpaired t test). CCh suppression of burst-evoked AHPs (**p<0.01 paired t tests) was unchanged between genotypes (mAHP ∼Δ1.5mV, sAHP ∼ Δ1mV, p>0.05 unpaired t test). **e** Example traces of isolated small-conductance Ca^2+^-activated SK currents and full inhibition by d-tubocurarine (dTC, grey). **f** dTC-sensitive current amplitude (left) and charge (right) for each genotype (WT n=8, HOM n=10 cells, 4 mice/genotype, p>0.05 unpaired t tests). **g** Example traces of spontaneous excitatory postsynaptic currents in WT and HOMS15A mice **h** Distribution of measured sEPSC properties for each genotype (WT: 1422-34 events, n=8 cells, HOM: 1455-65 events, n=7 cells; mean amp=14.0±0.8 *vs* 14.7±0.5 pA; instantaneous frequency=4.0±0.4 *vs* 4.3±0.3 Hz; N=3 mice/genotype; p>0.05, unpaired t tests).

**Supplementary Fig. 4:**
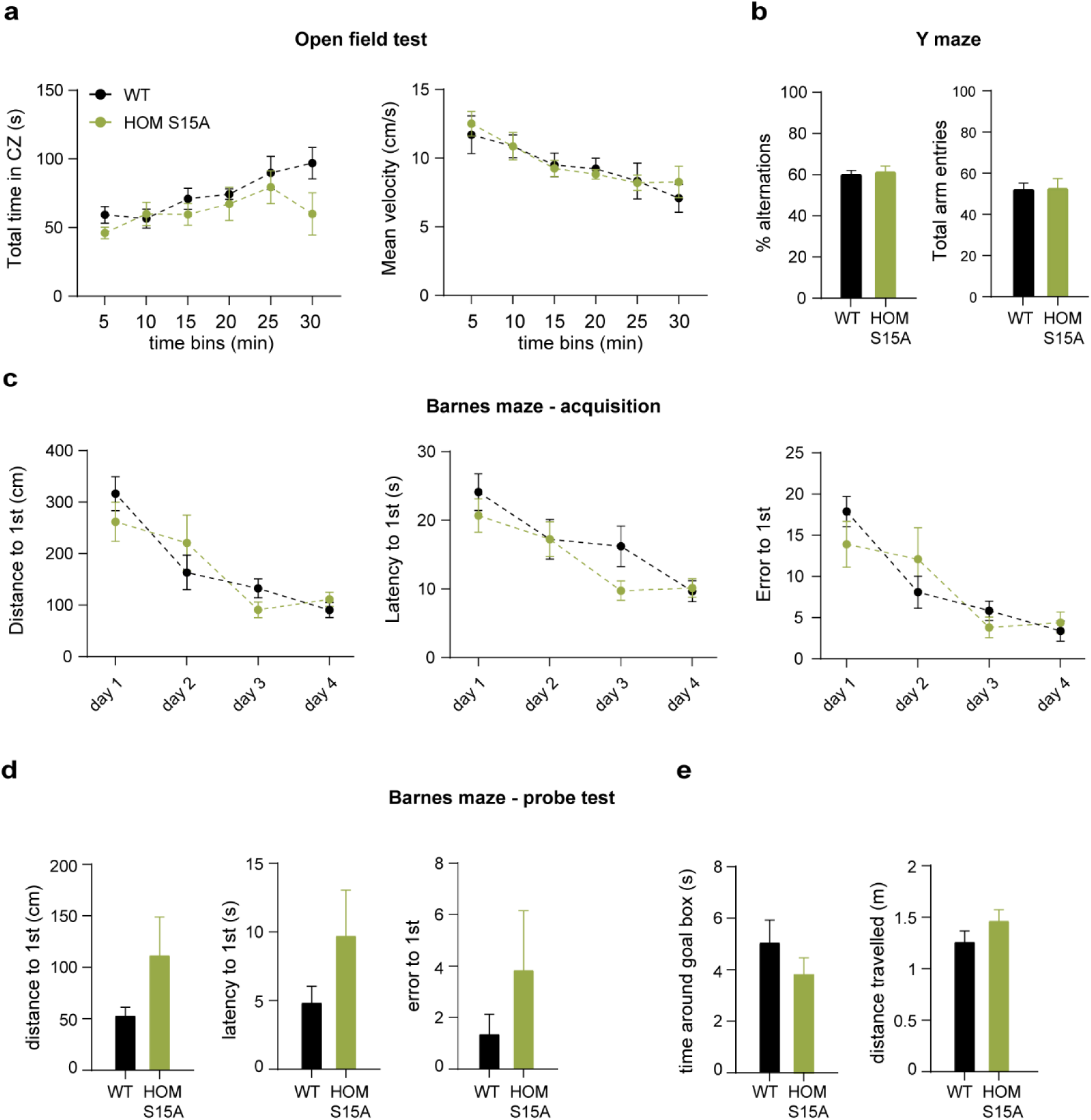
Behavioural characterization of Cav2.3 S15A phosphomutant mice. **a** left: total time in the central zone (CZ) of the open field arena in WT (N=8) and HomS15A (N=7) mice (30 min test) (p>0.05, Two-way Repeated Measures ANOVA; p=0.08 at 30 min, Fisher’s LSD); right: mean displacement velocity (p>0.05, Two-way Repeated Measures ANOVA). **b** Percentage of successful spontaneous alternations (left) and total arm entries (right) in the Y-maze test show no deficits in arm recognition and exploration in phosphomutants (p>0.05, unpaired t test). **c** Acquisition phase of the Barnes maze test with improved performance for both groups in the first 4 training days: distance travelled (left), latency (middle) and number of errors committed (right) during the 1^st^ visit to the escape box were not significantly different (p>0.05, Two-way Repeated measures ANOVA). **d** memory assessment on probe day 5 compared using the same parameters as in **c** (p>0.05, unpaired t tests). **e** Quantification of total time spent around the expected location of the goal box (left) and total distance travelled (right) (p>0.05 unpaired t tests).

